# Breathwork-Induced Psychedelic Experiences Modulate Neural Dynamics

**DOI:** 10.1101/2024.02.19.580985

**Authors:** Evan Lewis-Healey, Enzo Tagliazucchi, Andres Canales-Johnson, Tristan A. Bekinschtein

## Abstract

Breathwork is an understudied school of practices involving intentional respiratory modulation to induce an altered state of consciousness (ASC). We simultaneously investigate the phenomenological and neural dynamics of breathwork by combining Temporal Experience Tracing, a quantitative methodology that preserves the temporal dynamics of subjective experience, with low-density portable EEG devices. Fourteen novice participants completed a course of up to 28 breathwork sessions – of 20, 40 or 60 minutes – in 28 days, yielding a neurophenomenological dataset of 301 breathwork sessions. Using hypothesis-driven and data-driven approaches, we found that ‘psychedelic-like’ subjective experiences were associated with increased neural Lempel-Ziv complexity during breathwork. Exploratory analyses showed that the aperiodic exponent of the power spectral density – but not oscillatory alpha power – yielded similar neurophenomenological associations. Non-linear neural features, like complexity and the aperiodic exponent, neurally map both a multidimensional data-driven composite of positive experiences, and hypothesis-driven aspects of psychedelic-like experience states such as high bliss.

---

Mapping the relationship between subjective experience and the brain is a primary aim of the neuroscience of consciousness (Koch et al., 2016; Lutz & Thompson, 2003; Varela, 1996). Research over the last two decades has focused on identifying neural signatures associated with the breakdown of consciousness, in states such as dreamless sleep or anaesthesia. One overarching finding is that neural signal diversity, such as Lempel-Ziv (LZ) complexity, decreases in *states* of reduced consciousness (Schartner et al., 2015; Luppi et al., 2019; King et al., 2013; Pascovich et al., 2022). LZ complexity also has demonstrable utility probing the *contents* of consciousness (Canales-Johsnon et al., 2020; 2023), and as an index of arousal in patients with disorders of consciousness (Casali et al., 2013). However, when applied to altered states of consciousness (ASCs) that have multi-dimensional and rich subjective experiences, such as the ‘psychedelic state’ (Bayne & Carter, 2018; Fortier-Davy & Milliere, 2020), issues arise with this neuro-centric approach. Some studies have ignored the subjective experiences that occur from psychedelics (Barnett et al., 2020; Deco et al., 2018; Petri et al., 2014; Varley et al., 2020a), thus yielding neural signatures that may be broadly related to the *state* of consciousness that occurs, rather than the *contents* of consciousness itself.

To counter this, a group of researchers have recently emphasised the necessity to pair first-person subjective reports with neurophysiological data in the study of phenomenologically rich ASCs such as psychedelics, hypnosis, and meditation (Timmermann et al., 2023a). The methodology in question (neurophenomenology; Varela, 1996) is growing within the neuroscience of ASCs (Cardeña et al., 2013; Dor-Ziderman et al., 2013; Dor-Ziderman et al., 2016; Timmermann et al., 2019; Garrison et al., 2013; Berkovich-Ohana et al., 2013). Of note, Timmermann et al. (2019) conducted a neurophenomenological study of the psychedelic substance N,N-Dimethyltryptamine (DMT), finding associations between LZ complexity and the subjective intensity of the experience over time. Other research has found similar results, demonstrating that increased neural signal diversity is a robust signature of the ‘psychedelic state’ (Schartner et al., 2017; Mediano et al., 2024; Pallavicini et al., 2021)^1^, much like a breakdown of consciousness is associated with a reduction in neural signal diversity. According to the Entropic Brain Theory (EBT; Carhart-Harris et al., 2014; Carhart-Harris, 2018), this increase in neural informational complexity is associated with the heightened ‘richness’ of the psychedelic experience.

However, despite the increasing adoption of neurophenomenological approaches in consciousness science, we are still yet to fully understand how the specific *contents* of consciousness that arise in ASCs are related to neural signatures (Aamodt et al., 2021; 2023). This may be due to (i) a ‘uni-dimensional’ approach to studying the relationship between phenomenology and the brain (Bayne & Carter, 2018; Fortier-Davy & Milliere, 2020), (ii) the use of Likert scales to quantify the overall intensity of experiential dimensions, thus ignoring the temporal dynamics of phenomenology, and (iii) under-powered statistical tests in psychedelic studies.

We therefore present the current neurophenomenological research as a novel approach to the study of ASCs. To address aforementioned issue (i), we use data-driven techniques to reveal multi-dimensional phenomenological clusters (see Methods), which may be associated with neural signatures. To address issue (ii), we utilise temporal experience tracing (TET; Jachs et al., 2022; Jachs, 2021), a novel phenomenological methodology to empirically investigate subjective experience dynamics. TET involves the production of phenomenological timeseries, as participants retrospectively draw the intensity of pre-defined subjective experience dimensions over time on two-dimensional axes. By doing so, TET supersedes the capability of Likert scales, as these timeseries are more accurate reconstructions of subjective experience dynamics over time. Finally, to address issue (ii), we investigate high-ventilation breathwork, an ASC induced through voluntary changes in respiration (such as depth, breadth, and frequency; see Methods). The phenomenology of breathwork has been found to be similar to serotonergic psychedelics such as LSD and psilocybin (Puente et al., 2014; Bahi et al., 2023; Rock et al., 2015; Fincham et al., 2023a). Therefore, the study of this non-pharmacological ASC that is accessible to novices^2^, using TET and portable EEG devices, has yielded this neurophenomenological dataset, which is an order of magnitude larger than comparable psychedelic studies (e.g., Timmermann et al., 2019; 2023b; Schartner et al., 2017).

Breathwork encompasses a variety of distinct practices. For example, slow-paced breathing techniques induce a range of psychological outcomes, such as reduced arousal and enhanced relaxation (Zaccaro et al., 2018). However, within this study, we specifically investigate high-ventilation breathwork (Fincham et al., 2023a), in which participants couple a faster rate of breathing with breath retention phases to induce a hypoxic state (low blood oxygen saturation). Inducing changes in physiology through the modulation of the breath is one of the fundamental goals of high-ventilation breathwork (Kox et al., 2014), with the magnitude of these physiological changes linked to the intensity of the altered state of consciousness (Havenith et al., 2024).

Prior to the analysis of the data, neurophenomenological hypotheses were pre-registered on the Open Science Framework (https://osf.io/ef5j3). We focused initial analyses on LZ complexity, as previous research has found associations between psychedelic phenomenology and LZ complexity (Schartner et al., 2017; Timmermann et al., 2019; Mediano et al., 2024). The data were analysed using Data-Driven (DD), Hypothesis-Driven (HD), and Stimulus-Driven (SD) approaches, to elucidate the potential relationships between physiology, phenomenology, and neural complexity. Within the DD approach, we broadly hypothesised that k-means clustering (Likas et al., 2003) would extract a psychedelic-like phenomenological cluster, and this would be significantly different to other phenomenological clusters in terms of LZ complexity. Within the HD approach, we hypothesised that there would be an association between the retrospectively rated intensity of ‘bliss’ and LZ complexity. This was hypothesised as psychedelics robustly increase the phenomenological dimension of bliss (Hirschfield & Schmidt, 2021), which we theorised would load onto neural signatures associated with psychedelics, such as LZ complexity. Finally, within the SD approach, it was broadly hypothesised that breath retentions would modulate LZ complexity on a within-session basis (see Methods for a comprehensive overview).

## Methods

### Temporal Experience Tracing (TET)

TET has previously been used to study the phenomenological dynamics of different styles of meditation (Jachs et al., 2022; Jachs, 2022), is currently being used to study temporal dynamics of stress in autism (Gernert et al., 2023), has been applied to the study of chronic pain (Niedernhuber et al., 2023) and is being applied to two separate studies with the potent and fast-acting psychedelic DMT (publications ongoing). With the methodology of TET, dimensions of subjective experience most relevant to the phenomenon of study are identified. These dimensions are sub-components of the contents of consciousness, relating to affective, attentional, or motivational aspects of experience. After completing a session of the particular phenomenon of study, participants complete a retrospective trace of the intensity of each specific dimension.

At the inception of this study, there had been little research conducted on the subjective experiences of breathwork. Therefore, we sought to map out specific dimensions that form the phenomenological foundation of breathwork practices. SOMA Breath (https://www.somabreath.com/), an organisation that provides training and courses in breathwork practices, created a modified and streamlined version of an existing course for this study (under procedure), consisting primarily of breathwork sessions. The first author (ELH) completed this streamlined course. Each dimension was discussed with an experienced breathwork practitioner and with the CEO of SOMA Breath, and circulated to breathwork teachers for feedback. Fourteen dimensions were chosen as the most phenomenologically relevant for this particular breathwork modality. Five of these dimensions (marked with an asterisk) were also used in Jachs (2022). Below is a description of each dimension.

1. **Meta-awareness**\* is defined as being aware of the current contents of thought (Schooler et al., 2011). This has been highlighted as a fundamental construct in meditation practices, particularly open monitoring (Laukkonen & Slagter, 2021). This dimension was included as it may be a potential measure of the veridicality of the overall phenomenological report.
2. **Presence/Distraction** is defined as how present or distracted participants were during the breathwork session. An example provided to participants was, “…if you are not focused on putting effort into the breathwork session, then distraction would be particularly high at the moment. However, if you are ‘with’ the breathwork practice, being at one with your experience as it arises, you are highly present, and thus score low in distraction.”
3. **Physical Effort** refers to the amount of physical exertion participants exhibited during the breathwork experience. The breathwork sessions can be physically demanding, due to an increase in breathing frequency or sustained breath holds. It was therefore deemed a key dimension in the practice of breathwork across participants.
4. **Mental Effort**\* may be defined as the exertion of attentional resources under increasingly demanding task-conditions (Alnæs et al., 2014). Mental effort has also been contextualised within the terms of the costliness of cognitive control (Shenhav et al., 2017).
5. **Boredom**\* has received much attention in the scientific literature (Fisherl, 1993; Smith, 1981). The phenomenology of boredom has been characterised by feelings of restlessness, lethargy, lack of concentration, and distorted time perception (Martin et al., 2006). A review study has highlighted that boredom is related to an inability to sustain attentional capacities (Westgate & Steidle, 2020). It may therefore be related to the construct of mental effort.
6. **Receptivity/Resistance** refers to a state in which a participant is resistant to the experiences that occur within the breathwork practice. Being receptive to the successive experiences is the antonym of resistance, and may be phenomenologically similar to the state of ‘surrender’, which has been highlighted as a baseline predictor of positive reactions to psychedelic experiences (Aday et al., 2021).
7. **Emotional Intensity**\* was a dimension related to the intensity of their emotional state, regardless of positive or negative valence. This setup, of retrospectively rating the intensity of emotions in a continuous fashion, was also employed in a study using virtual reality (Hofmann et al., 2021). The researchers found that emotional intensity was associated with parieto-occipital alpha power, which was further confirmed through decoding analysis.
8. **Clarity**\* refers to the vividness of the contents of consciousness (Lutz et al., 2015). In the case of perceptual clarity, we may refer to visual vividness as analogous to this. In previous studies, endogenous attention has demonstrably increased the perceived contrast of stimuli in a rapid serial visual presentation task (Liu et al., 2009). While visual vividness may be included in the experience of clarity, other aspects of conscious experience are also perceived with a degree of clarity, such as affective tone.
9. **Release/Tension** refers to the amount of either physical, or psychological tension that is experienced. It is hypothesised that high periods of tension may be followed by high periods of release. There may be some overlap with these periods of high intensities of release and “emotional breakthrough” (Roseman et al., 2019), which has its roots in the psychoanalytic concept of catharsis (Breuer & Freud, 1895; Jackson, 1994). Emotional breakthrough may represent a large aspect of the phenomenology of the psychedelic experience (Carhart-Harris et al., 2021), and may be a potential therapeutic mechanism.
10. **(Dis)embodiment** refers to the experience of not identifying with one’s own body (Blanke, 2012). The antonym of this, the feeling of identifying with and normally perceiving one’s body, has frequently been investigated and modulated in studies on self-consciousness (Blanke & Metzinger, 2009). Disembodiment forms a specific sub-component of the 11 Dimensions of Altered States of Consciousness questionnaire (11D-ASC; Studerus et al., 2010), and has been modulated through the administration of psychedelics (Bernasconi et al., 2014; Kometer et al., 2012; Kraehenmann et al., 2017). Bahi et al. (2023) found that a conscious connected breathwork session increased feelings of disembodiment similar to a psilocybin dose of 400μg/kg. Within the present study, this dimension was coded so high values denoted a high intensity of embodiment, and low-values denoted a high intensity of disembodiment.
11. **Bliss** is another sub-component of the 11D-ASC (Studerus et al., 2010). It may be defined as experiencing boundless pleasure, or a profound peace. The empathogen/entactogen 3,4-methylenedioxy-methamphetamine (MDMA) has been found to significantly increase a blissful state (Studerus et al., 2010). Again, Bahi et al. (2023) found that a conscious connected breathwork session yielded similar ratings of bliss when compared to 400μg/kg of psilocybin.
12. **Insightfulness** was also taken from the 11D-ASC (Studerus et al., 2010). Within the literature, insight may refer to the experiences of eureka moments. These are characterised as sudden moments of clarity into the solution of a problem (Kounios & Beeman, 2014). Cognitive (neuro)science has attempted to identify the neural correlates of the eureka moment (Aziz-Zadeh et al., 2009; Tik et al., 2018), and assessed the effect of artificially induced eureka moments on metacognition (Laukkonen et al., 2020). However, as eureka moments are temporally localised phenomena, this specific dimension is focussed on assessing the novelty of the stream of consciousness. As above, Bahi et al. (2023) found that retrospective ratings of Insightfulness were similar to 400μg/kg of psilocybin.
13. **Anxiety** was defined as the experience of dysphoria, characterised by a strong, negatively valenced state. Within studies that evaluated the subjective effects of psychedelic substances, the experience of anxiety has been associated with a lack of response for ketamine-assisted therapy for major depression (Aust et al., 2019). Anxiety also forms a sub-component of the 11D-ASC (Studerus et al., 2010).
14. **Spiritual Experience** is the final dimension included from the 11D-ASC (Studerus et al., 2010). This dimension is qualitatively similar to peak (Stace, 1960), or mystical (Hood, 1975; Griffiths et al., 2006; Barrett & Griffiths, 2018) experiences, characterised by feelings of intense awe (Hendricks, 2018), and feeling connected to a higher power (Studerus et al., 2010). The mystical experience forms part of psychedelic phenomenology at higher doses, and may be a predictor of clinical benefits of psychedelics (for a review, see Johnson et al., 2019).

### EEG Headsets

Participants were provided with a portable low-density Dreem EEG headset (Arnal et al., 2020). The EEG headsets collect both neural and accelerometry data. These headsets have seven channels. Each channel is a longitudinal bipolar montage between frontal electrodes (F7, F8, and FPz), or frontal and occipital electrodes (O1 and O2). The sampling rate of the headset was 250Hz, and the resolution of the AD converter was 24 bits. The hardware did not apply any filtering or preprocessing of the data. One channel was re-referenced to provide solely occipital-occipital information.

### Participants

The experimental protocol was approved by the Cambridge Psychology Research Ethics Committee. All participants took the full breathwork course free of charge and were not directly reimbursed for their time.

Participants were recruited via the SOMA Breath advertising channels on social media. The eligibility criteria was the following:

● Between 18-65 years of age
● Participated in two or less sessions of high-ventilation breathwork practices
● Based in the UK and able to complete the whole study
● No chronic mental health, sleep-related or neurological medical problems.

21 participants were originally recruited for this study. All participants signed an informed consent form. One participant withdrew due to pregnancy, three participants withdrew due to unrelated health issues, one participant withdrew due to a bereavement, and one participant withdrew due to a lack of time in the mornings to complete the sessions. Finally, one participant returned the physical TET data in a damaged condition, and thus was excluded from further analysis. Therefore, data is presented from 14 participants (10f, 4m; average age = 40, SD = 9).

### Procedure

Participants completed a four-week course delivered online via SOMA Breath. As mentioned, SOMA Breath is a company that provides training and facilitation of breathwork sessions. The four-week course was delivered remotely online. Within this course, participants completed a breathwork session every day for 28 days. The breathwork session is an online audio track, one for each corresponding week of the course. The tracks themselves are composed of breathing instructions, and are accompanied by evocative background music. As the course progressed, the sessions became longer, with more breath holds, inducing a more pronounced acute hypoxic state (Kox et al., 2014).

Prior to the start of the course, participants were provided with a personal low-density EEG device, and either a TET booklet or digital folder of powerpoint presentations. The booklet and powerpoint presentations contained TET graphs for each breathwork session, with markers along the x-axis corresponding to the conditions within the session (see Supplementary Figure S1 for an example). In addition to this, the participants met with the primary researcher to (i) learn how to perform the tracing method, (ii) learn how to use the portable EEG headset, and (iii) discuss the definition of each phenomenological dimension.

As previously mentioned, the breathwork course was organised into four separate weeks, with the same breathwork session practised each day of the week. For the first and fourth week of the course, the breathwork session was approximately twenty minutes long. For the second week, the session was approximately forty minutes long. For the third week, the session was approximately one hour long. On the first day of each week of the course, the participants met online with the CEO of SOMA Breath, where the breathwork technique was taught, questions could be answered, and the breathwork session for that week was conducted. Succeeding this, each participant was to complete the breathwork sessions daily, yet of their own accord (with recommendation to practise it in the morning). Each session was partitioned into five different instructions (hereby named conditions):

1. **Introduction**: Comprises the first part of the session, where there are no specific instructions and only music is played.
2. **Breath**: Comprises instructions to breathe in a rhythmic and standardised manner.
3. **Hold**: Within this condition, the participant is instructed to actively hold their breath to induce a state of acute mild hypoxia. In some cases, the participant is instructed when to stop holding the breath, but in the later weeks of the course, there is less explicit instruction as when to conduct this.
4. **Fast**: This section applies to the third and fourth weeks of the course. Within these sections, the participant is instructed to breathe at a rate of one inhalation/exhalation every two seconds.
5. **End**: This represents the end of the session, where there is no instruction given, and only music is played.

Participants were instructed to complete the daily breathwork sessions with eyes closed. Participants wore the EEG headset during these sessions. Succeeding each session, participants completed TET graphs for each of the fourteen phenomenological dimensions. Participants also provided qualitative data regarding subjective experiences during the corresponding breathwork session. They conducted this by using voice-recording software on Microsoft Word, filling in a personalised sheet corresponding to each session. A schematic overview of the course and subsequent analyses can be found in Figure 1.

**Figure 1:**
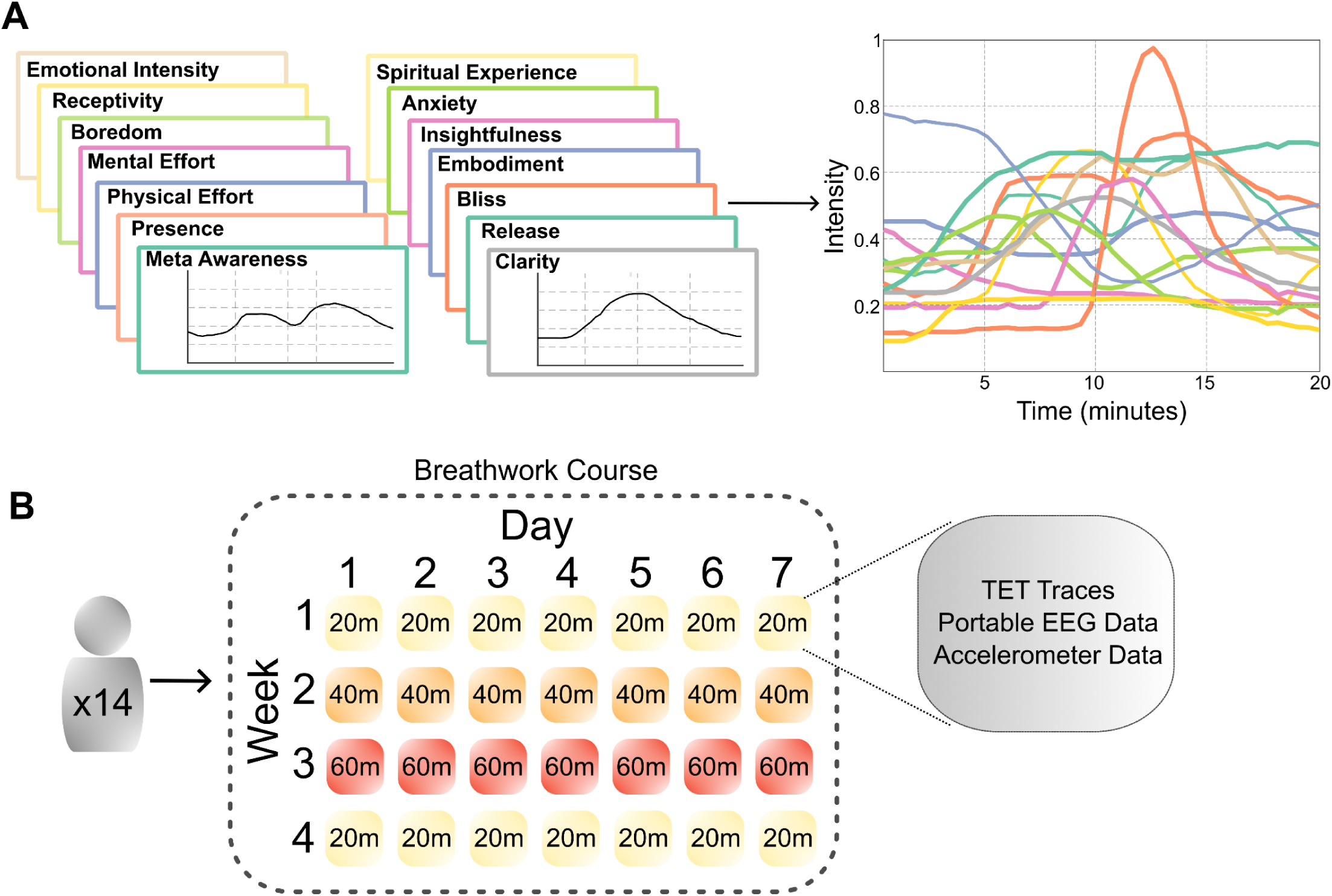
A schematic overview of Temporal Experience Tracing (TET), and the breathwork course. A) After a breathwork session, a participant will retrospectively trace the felt intensity for each distinct phenomenological dimension over time (left). These traces are then digitised into vectors and concatenated, thus forming a 14-dimensional matrix of phenomenological intensity (right). B) 14 participants completed a 28-day breathwork course, organised into four separate weeks. Within each week, the participant conducts the same breathwork session every day for seven days. Between weeks, the sessions become more intense; they increase in length, and comprise more breath holds within the session, allegedly culminating in more intense hypoxic states. During each session, participants recorded EEG and accelerometry data using the Dreem EEG headband (Arnal et al., 2020). After each session, the participants reported the subjective intensity of 14 distinct phenomenological dimensions over time. TET data was preprocessed and organised into experience clusters using k-means. LZ complexity measures were computed on the EEG data, as pre-registered. Exploratory analyses were conducted on the oscillatory power and aperiodic exponent of the EEG signal.

In addition to the above, 7 participants completed a phenomenological interview regarding their subjective experiences of note during the breathwork course. The phenomenological interview lasted up to two hours. These findings are partial and not described in this report.

### Preprocessing **-** TET Data

TET data were converted to vectors using a semi-automatic custom MATLAB script. The script converted the image coordinates of the TET data to graph coordinates, based on the orientation of the x and y axes. Each phenomenological dimension was manually checked to ensure the vectors were similar to the TET traces themselves. After conversion, each phenomenological dimension was concatenated in a column-wise fashion, creating a 14xN TET matrix for each breathwork session, where N is the corresponding number of data points generated for that session. Each participants’ TET data were normalised across all of their sessions for the whole course. Based on the assumptions of previous datasets (Jachs, 2022), each datapoint represented 28 seconds of the breathwork session. Sessions missing one or more dimensions were excluded from further analysis.

### Preprocessing – Neural and Accelerometer Data

Low-pass filtered (35Hz) neural data was downloaded from the Dreem server after the session was automatically uploaded. As impedance could not be assessed with the Dreem headsets, the data quality was assessed during the preprocessing phase. Neural data was high-pass filtered at 1Hz, and cut into four-second epochs. Within the course, participants were instructed to tap on the headset three times at the beginning of each breathwork session. Each dataset was manually scanned to identify this within the signal, and the session was cut to this corresponding starting point. Each epoch was labelled according to one of the five aforementioned conditions (i.e. according to ‘Introduction’, ‘Breath’, ‘Hold’, ‘Fast’, or ‘End’) based on the instructions of that session. The data was cleaned using a custom script designed to automatically reject epochs exceeding ±300μv. For the hypothesis-driven approach, using association metrics between the intensity of bliss and neural complexity, sessions with more than 50% of the neural epochs rejected were excluded. For the data-driven approach, and the stimulus-driven approach, sessions with over 80% of the neural epochs rejected were excluded. Further to this, consistent noisy channels at the participant level which led to the rejection of all epochs were excluded in order to keep these sessions for further analysis. Subjects 11 and 19 had two noisy frontal paired channels, which were excluded from further analysis. For each session, breathing frequency was extracted from the z-axis of the accelerometer on the EEG headband. The data were bandpass filtered between 0.02 and 0.2Hz and detrended.

## Analyses

### Lempel-Ziv Complexity

We focused our preregistered hypotheses on LZ complexity measures, as it is a relatively simple method that has been robustly applied to cognitive neuroscientific work on consciousness (Casali et al., 2013, Frohlich et al., 2022; Pascovich et al., 2022; Schartner et al., 2015; 2017; Timmermann et al., 2019). Briefly, LZ complexity is computed through the binarization of the signal around its median, where all values above the signal are 1, and all values below the signal are 0. The signal is then algorithmically scanned sequentially for novel patterns within the signal (Kaspar & Schuster, 1987), creating a ‘dictionary’ of binary sequences for each timeseries. The length of this dictionary is a measure of the complexity of the signal, which can then be standardised through the division of this by the computed complexity of the same data that is randomly shuffled.

For the current study, LZ complexity was computed in two forms: concatenated LZ (LZc) and summed LZ (LZSum). For LZc, all channels are concatenated together to form one timeseries, with the computations above applied to the single concatenated signal. In the second form, the LZ computation above was applied to each channel separately, with the results averaged. We applied these two computations to investigate potentially discrepant results, as observed in previous research (Schartner et al., 2015).

The above computations are a measure of global signal complexity. However, both LZSum and LZc were computed on the separate channel pairing groups (frontal with frontal, frontal with occipital, and occipital with occipital), to investigate spatial contributions to complexity changes within the varying hypotheses. The variations in the computation of LZ complexity, based on spatial contribution and LZSum/LZc, are hereby referred to as complexity types.

### Phenomenological Clustering

In order to extract metastable clusters of subjective experience that occurred throughout the breathwork course, we applied k-means clustering (Likas et al., 2023) to the TET data, using squared Euclidean distance as the distance metric. This is a data-driven analysis approach, as the clustering algorithm receives no *a priori* information regarding the structure of the phenomenological data, or about which subject, week, or session, the phenomenological data belongs to.

The optimal clustering solution was found through the Calinski-Harabasz criterion (Calinski & Harabasz, 1974). Furthermore, as the clustering solution depends on the initiation point of the centroids, the algorithm was replicated 1000 times with randomly chosen initiation points. The resulting clusters with the lowest sum of squared distances were subsequently chosen.

### Stimulus-Driven Analyses

The z-axis accelerometer data was extracted from the Dreem EEG device. The data were filtered between 0.02 and 0.2Hz, detrended, and normalised to have zero mean and a standard deviation of one. The data was cut to the length of the session, and the ‘Hold’ sections of the session were identified. Within these specific windows, the *findchangepts* function in MATLAB was used to identify the most abrupt change in the signal. This timepoint was an approximation of when the participant took their first inhale after holding their breath for a certain amount of time. The LZ complexity of this neural epoch (at the temporal scale of 28s) was compared to the neural epoch prior to the start of the breath hold.

We identified these epochs as we aimed to use the accelerometer data as a proxy for the hypoxic state of an individual (as the headsets lacked pulse oximeter data). The motivation behind the identification of the first breath was based on previous research on intermittent hypoxia-based breathwork practices showing that blood oxygen concentration is lowest at the end of the breath hold sections (Kox et al., 2014). We therefore aimed to investigate whether these sections related to changes in neural complexity at the session-level.

### Statistical Analyses

All statistical analyses were carried out with MATLAB R2022a (The Mathworks Inc., 2022). For each analysis approach, linear mixed models were computed. In addition to this, generalised linear mixed models with a Logit link function were computed for the exploratory data-driven approach (see below), to input the neural features into the same model, investigating whether each variable may ‘explain away’ the effects of LZ complexity. In order to statistically model the experimental design of the study, we included subject, week, and session as nested random effects for each model. Results below are presented on data that has been centred and cleaned of outliers above and below 2.5 standard deviations of the mean LZ complexity. Further, to compare the outputs of the mixed models for the variety of LZ spatial complexity types, each LZ complexity variable was standardised so that the standard deviation was 1. FDR multiple comparisons were computed to adjust the p-values of the models (Benjamini & Hochberg, 1995) using the *mulitcmp* function in MATLAB (Penn, 2023).

### Exploratory Analyses

To further investigate other neural signatures associated with psychedelics and subjective experience states, we explored the relationship between aperiodic and periodic components of the EEG power spectral density (PSD) and phenomenology. We used the fitting oscillations and one-over-f toolbox (fooof; Donoghue et al., 2021), to model periodic and aperiodic components of the power spectra. The fooof toolbox models the offset and exponent of the neural power spectra, and identifies peaks in the signal corresponding to oscillatory power. This therefore distinguishes between oscillatory power and aperiodic components of the EEG PSD.

Peak frequencies were identified and categorised within the canonical frequency bands delta (1-4Hz), theta (4-8Hz), alpha (8-13Hz), and beta (13-30Hz). Gamma oscillations were excluded from this analysis as the data was low-pass filtered at 35HZ, and this frequency band may more likely reflect muscular activity in EEG (Muthukumaraswamy, 2013). Much like the LZ computation above, the relative oscillatory power was subdivided into spatial areas of the EEG device (Frontal with Frontal, Frontal with Occipital, Occipital with Occipital, and Global). The above computation was applied on each four second neural epoch. To bridge the gap between phenomenological data and oscillatory power, the median value of the components taken for every seven neural epochs (equating to one phenomenological datapoint at 28 seconds) were input for further statistical analyses. If there were multiple oscillatory peaks within the same frequency band in the same neural epoch, the peak power was summed together^3^. While we conducted exploratory analyses for all frequency bands, we present results primarily for alpha power within this paper, as a reduction in alpha oscillatory power is another robust neural signature of the psychedelic state (Valle et al., 2016; Timmermann et al., 2019; Pallavicini et al., 2019; Mediano et al., 2024).

We also explored the relationship between the aperiodic exponent of the neural power spectral density (PSD) and phenomenology. We intended to investigate the aperiodic exponent as psychedelics have a demonstrable impact at the broadband level (Muthukumaraswamy et al., 2013), and measuring the aperiodic exponent may tap in to physiological mechanisms that may be associated with breathwork (such as excitation/inhibition balance; Gao et al., 2017; Fincham et al., 2023a).

## Results

In total, 301 breathwork sessions were included for phenomenological clustering (Week 1 = 88, Week 2 = 79, Week 3 = 73, Week 4 = 61). The hypothesis-driven statistical analyses were conducted on 146 sessions (Week 1 = 37, Week 2 = 49, Week 3 = 34, Week 4 = 26), data-driven statistical analyses were conducted on 204 sessions (Week 1 = 62, Week 2 = 60, Week 3 = 45, Week 4 = 37), and stimulus-driven statistical analyses were conducted on 201 sessions (Week 1 = 61, Week 2 = 59, Week 3 = 44, Week 4 = 37). For an overview of the number of sessions each subject contributed to the analysis, see Supplementary Figures S2-S5. It is worth noting that, due to the more conservative session rejection threshold (if more than 50% of the EEG epochs were rejected) combined with fewer Week 4 sessions completed, only 9/14 participants contributed Week 4 sessions to the analysis, with two of these contributing nearly half of all sessions.

## Data-Driven

### K-means Extracted Two Phenomenological Clusters: One Loading onto Negative Physical and Mental Dimensions, the Other on ‘Psychedelic-Like’ Dimensions

K-means clustering was applied to the TET data for all breathwork sessions. The Calinski-Harabasz criterion (Calinski & Harabasz, 1974) found the optimal number of clusters to be two. Clusters were of approximately equal size (12,460 data points in cluster one; 11,660 data points in cluster two). Linear mixed models were computed to assess the difference between clusters on each phenomenological dimension. With the exception of Embodiment, all phenomenological dimensions were significantly different between clusters (Figure 3A. Cluster two was significantly lower in the following dimensions (compared to cluster one): Meta-Awareness (–0.06, SE=0.015, t(24,118)=-4.16, p<0.001), Presence (–0.25, SE=0.011, t(24,118) = –21.58, p<0.001), Receptivity (–0.2, t(24,118)=-15.72, SE=0.012, p<0.001), Emotional Intensity (–0.14, SE=0.012, t(24,118)=-12.26, p<0.001), Clarity (–0.18, SE=0.012, t(24,118)=-15.66, p<0.001), Release (–0.21, SE=0.012, t(24,118)=-18.18, p<0.001), Bliss (–0.24, SE=0.01, t(24,118)=-23.15, p<0.001), Insightfulness (–0.16, SE=0.01 t(24,118)=-14.64, p<0.001), and Spiritual Experience (–0.2, SE=0.01, t(24,118)=-19.6, p<0.001). Cluster two was significantly higher in Physical Effort (0.07, SE=0.013, t(24,118)=5.28, p<0.001), Mental Effort (0.15, SE=0.012, t(24,118)=11.43, p<0.001), Boredom (0.17, SE=0.013, t(24,118)=13.77, p<0.001), and Anxiety (0.09, SE=0.001, t(24,118)=9.85, p<0.001) (see Supplementary Table S1 for outputs of each mixed model). It is of note that bliss, insightfulness, and spiritual experience are all sub-components of Oceanic Boundlessness, which are strongly associated with the psychedelic experience (Roseman et al., 2018). Comparing these clusters therefore represents a multivariate and theory independent way to examine neural differences between (quantitative) phenomenological dimensions most associated with the psychedelic state.

**Figure 2:**
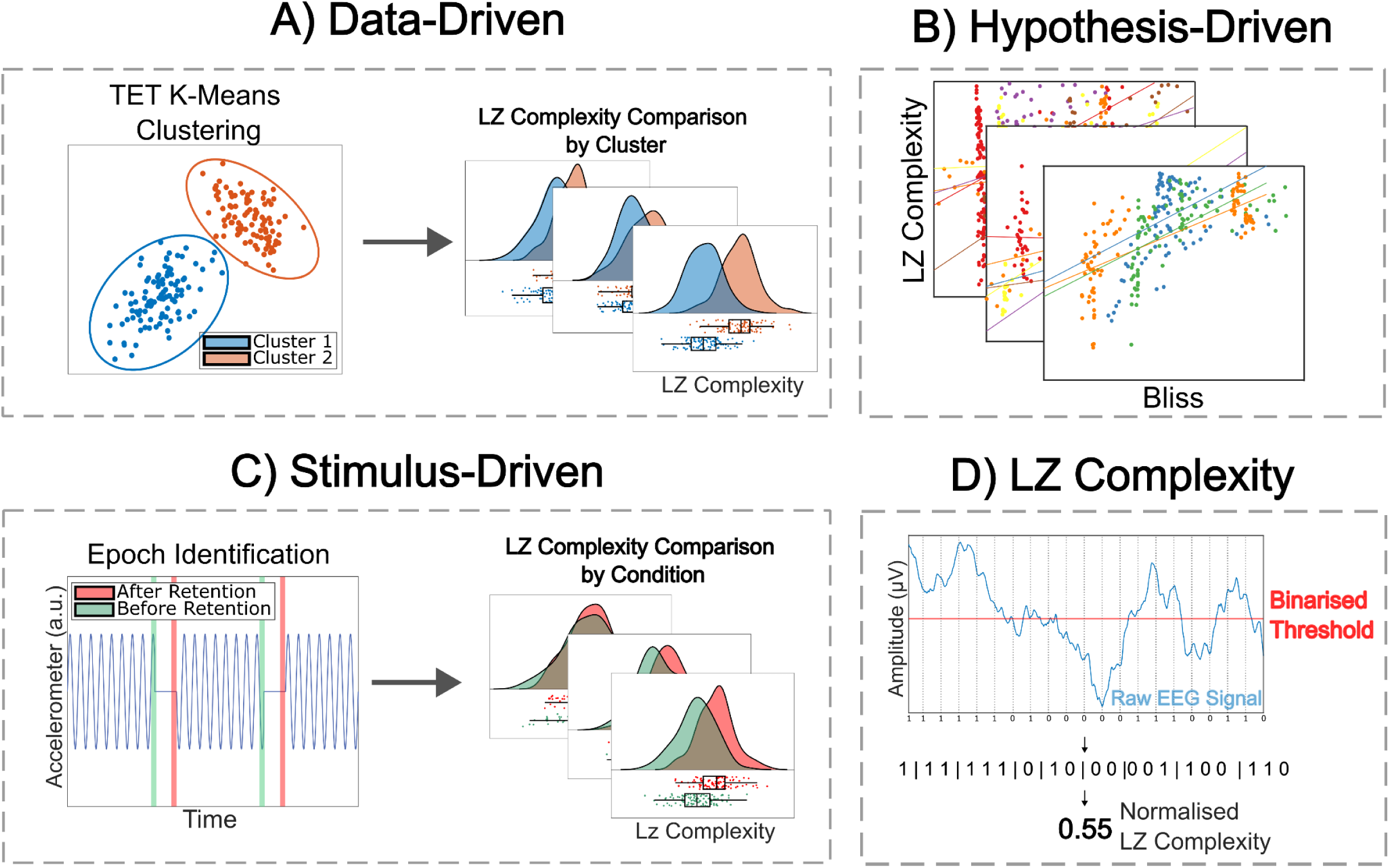
A schematic overview of the three analysis approaches. A) K-means clustering is applied to the TET data from all sessions, and linear mixed models are used to investigate the difference in LZ complexity between clusters, whilst factoring in the random variance of each session. B) Linear mixed models are used to compute the association between LZ complexity and the TET dimension ‘bliss’. Each colour within one panel represents the bliss and LZ complexity data for one session within one week for one participant. The linear mixed models are based on evaluating the differing slopes for each individual session, thus factoring in the random variance of each subject, week, and session.. C) Epochs in which the participant took their first breath during the breath hold sections are identified using accelerometer data, and compared to an opposing epoch in a session-wise fashion (again using linear mixed models). D) A schematic illustrating the computation of LZ complexity. For an exemplary epoch of EEG data, the raw signal is converted into a binary score for whether each data point is above or below the binarised threshold. This binarised timeseries is then expressed through the length of a ‘dictionary’, in which a new dictionary entry refers to a novel sequence within the timeseries. The length of the dictionary is the LZ score – analogous to the compressibility of the signal – which can be normalised by dividing this by the LZ score of the randomly shuffled epoch.

**Figure 3:**
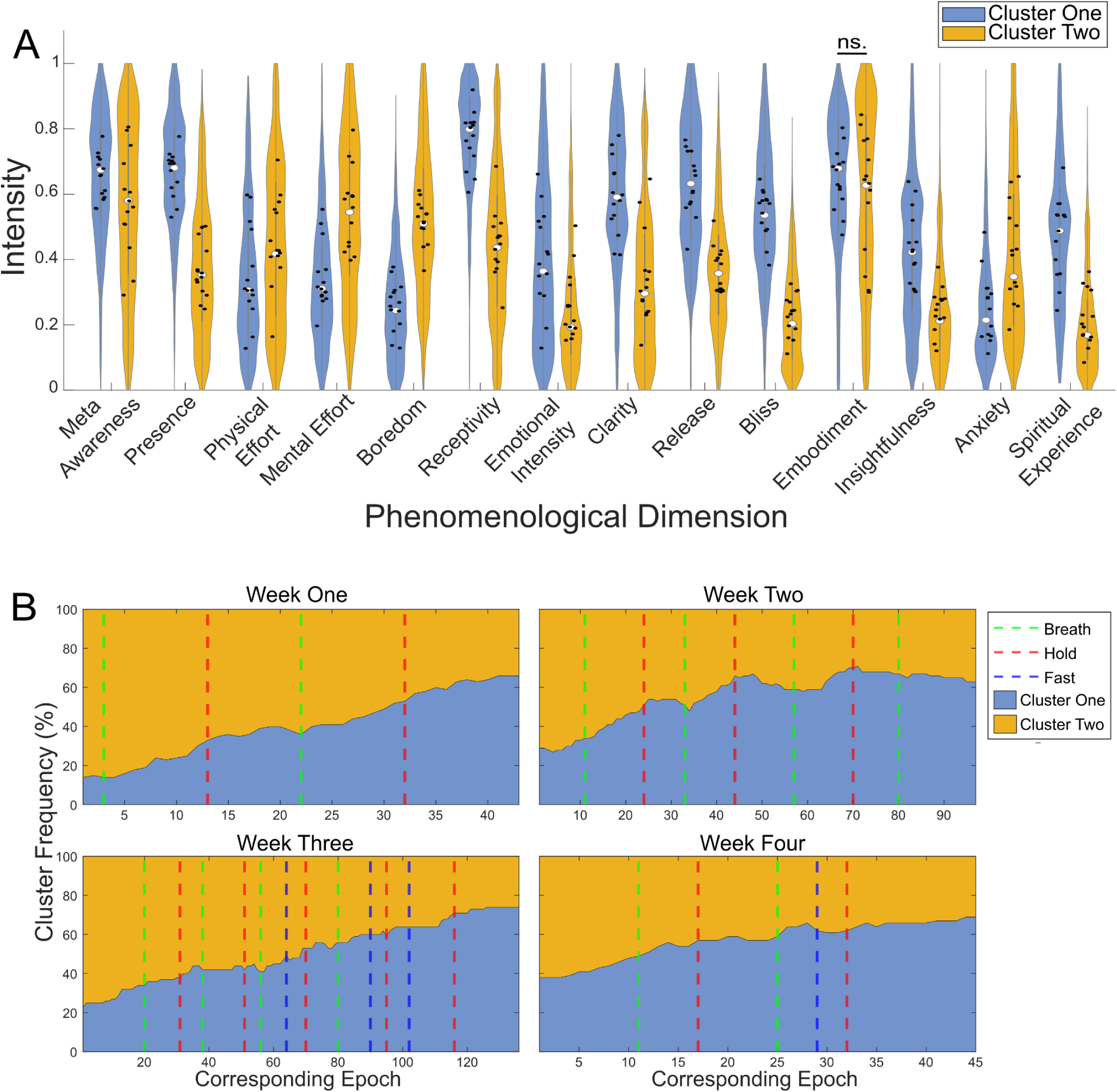
The breathwork session intensity (length) increased the likelihood of the ‘psychedelic-like’ cluster. A) Violin plots showcasing the data distributions for each phenomenological dimension per cluster. Excluding ‘Embodiment’, each phenomenological dimension was reliably different between clusters. Black filled circles are the mean intensity for each subject. B) Sandbox plots representing the proportion of time spent in each phenomenological cluster by week. Participants were ∼1.5x more likely to be in cluster one in week three when compared to week one. However, this effect was not driven by within-session changes.

### The Breathwork Session Intensity (Length) Increased the Likelihood of the ‘Psychedelic-Like’ cluster

Based on the idea that phenomenology would be associated with the intensity of the breathwork session conducted (i.e., a dose-response relationship), we hypothesised that cluster one was more likely to appear in the more intense breathwork sessions (i.e., weeks two and three). When comparing weeks one and four with weeks two and three, the likelihood odds ratio (OR) that cluster one was more likely to occur than cluster two was 1.27 (95% CI [1.20, 1.34]). When comparing week one to week three, the OR that cluster one was more likely to occur than cluster two was 1.54 (95% CI [1.43, 1.67]). To investigate whether this effect held at the within-session level, we also computed the within-session odds ratio, and investigated whether these significantly differed by week. The odds ratio was computed by dividing the amount of Cluster 1 epochs in the second half of the session by the amount of Cluster 1 epochs in the first half of the session. We then log transformed these values due to the exponential nature of odds ratios in this situation. A larger odds ratio would indicate a higher likelihood that Cluster 1 is found at the end of the session compared to the first. After excluding infinite values, we ran a linear mixed model with the formula ‘Log_Odds_ratio ∼ Week + (1 | Subject)’. When comparing the odds ratio to Week 1, we found no significant differences in Week 2 (0.05, SE = 0.14, t(163) = 0.36, p = 0.72), Week 3 (0.062, SE = 0.14, t(163) = 0.42, p = 0.67), or Week 4 (0.093, SE = 0.15, t(163) = 0.61, p = 0.54). This suggests that, while the cluster associated with psychedelic phenomenology may be contingent on the intensity of the breathwork session conducted (Figure 3B), these may not be driven by changes within-session.

### Psychedelic-Like Phenomenological Cluster is Higher in Lempel-Ziv Complexity

To assess the difference in LZ complexity between phenomenological clusters, we computed linear mixed models for each spatial LZ complexity type, with Cluster as the predictor variable. In order to balance parsimony with complexity of the model, we used subject, week, and session as nested random factors, with week and complexity type as fixed effects. Due to the high degree of correlation between the LZ complexity types (max correlation = 0.89), separate linear mixed models were computed for each spatial complexity type. All spatial complexity types showed that cluster two was significantly lower in complexity compared to cluster one: Frontal – Occipital (F-O) LZSum (–0.2, SE=0.044, t(13,925)=-4.47, p<0.001), Frontal – Frontal (F-F) LZSum (–0.19, SE=0.046, t(13,972)=-4.07, p<0.001), Occipital – Occipital (O-O) LZSum (–0.23, SE=0.043, t(14,117)=-5.39, p<0.001), Global LZSum (–0.25, SE=0.048, t(13,868)=-5.18, p<0.001), F-O LZc (–0.22, SE=0.046, t(13,882)=-4.42, p<0.001), F-F LZc (–0.15, SE=0.046, t(13,975)=-3.3, p<0.001), and Global LZc (–0.16, SE=0.043, t(13,724)=-3.7, p<0.001). The average session-level difference between clusters by week is visualised in Figure 4B.

**Figure 4:**
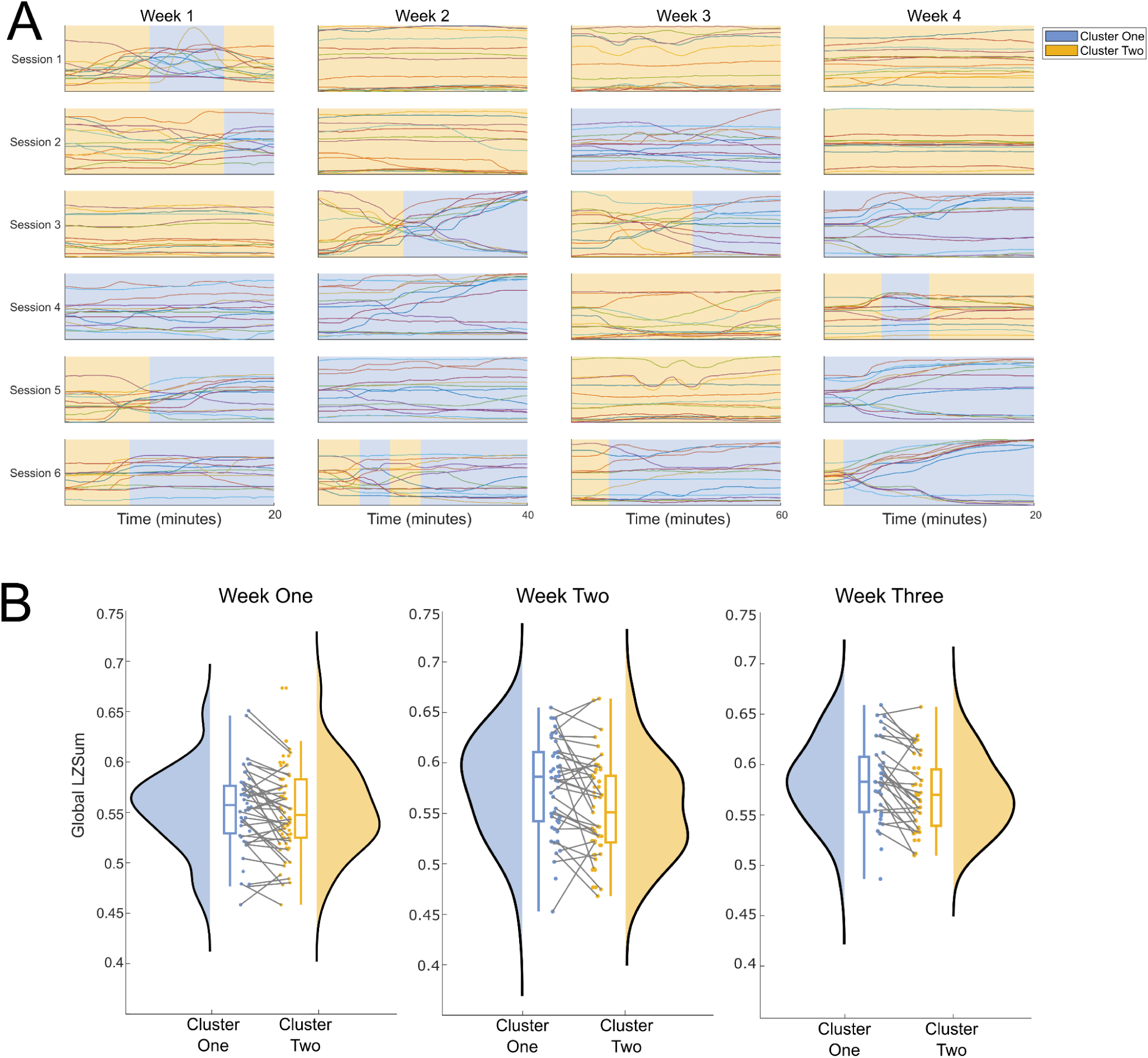
The psychedelic-like phenomenological cluster is reliably higher in LZ complexity across weeks. A) An example of the clustering algorithm as applied to each session for one participant. Each plot contains the TET data for each of the fourteen phenomenological dimensions, with the corresponding cluster identity shaded in blue or yellow for cluster one and cluster two respectively. B) Raincloud plots depicting the average Global LZSum values in a session-wise fashion for each phenomenological cluster show reliably higher complexity for cluster one compared to cluster two in every week. Each data point is the average complexity value for each cluster within one session, with connecting lines depicting the complexity difference between clusters yet within sessions. For parsimony, only weeks 1-3 are shown here. See Supplementary Figure S6 for week four.

### Hypothesis-Driven

Within the hypothesis-driven approach, the association between the retrospectively rated intensity of bliss (as a proxy psychedelic-like phenomenological dimension) and neural complexity was assessed.

### Bliss is Associated with Neural Complexity but not Modulated by the Intensity of the Breathwork Session

We first hypothesised that there would be an association between bliss and neural complexity, as the phenomenological dimension of bliss increases as a function of psychedelic use (Hirschfield & Schmidt, 2021; Roseman et al. 2018). To assess this potential relationship, we used linear mixed models, with week and complexity type as fixed effects, and subject, week, and session as nested random effects. All complexity types were significantly associated with the phenomenological dimension of bliss, where an increase in bliss was associated with an increase in complexity at the population level: F-O LZSum (0.04, SE=0.006, t(11,298)=6.24, p<0.001), F-F LZSum (0.03, SE=0.008, t(11,350)=4.06, p<0.001), O-O LZSum (0.06, SE=0.007, t(11,489)=8.61, p<0.001), Global LZSum (0.04, SE=0.007, t(11,248)=5.76, p<0.001), F-O LZc (0.04, SE=0.005, t(11,324)=7.02, p<0.001), F-F LZc (0.03, SE=0.008, t(11,349)=4.26, p<0.001), and Global LZc (0.05, SE=0.007, t(11,124)=6.81, p<0.001).

Additionally, we hypothesised that the magnitude of association would increase as a function of the length of the breathwork session conducted. We tested this hypothesis through the inclusion of week as an interaction term. For these models, all main effects remained significant, where an increase in bliss was associated with an increase in complexity at the population level: F-O LZSum (0.04, SE=0.013, t(11,295)=3.24, p=0.004), F-F LZSum (0.04, SE=0.016, t(11,347)=2.62, p=0.02), O-O LZSum (0.06, SE=0.014, t(11,486)=4.22, p<0.001), Global LZSum (0.04, SE=0.014, t(11,245)=2.45, p=0.035), F-O LZc (0.05, SE=0.01, t(11,321)=4.84, p<0.001), F-F LZc (0.04, SE=0.016, t(11,346)=2.68, p=0.02), and Global LZc (0.05, SE=0.02, t(11,121)=3.81, p<0.001). However, no interaction effects were significant when comparing the magnitude of association to the first week (Supplementary Figure S7). This suggests that the association between bliss and neural complexity is relatively stable across weeks of the breathwork course (Figure 5).

**Figure 5:**
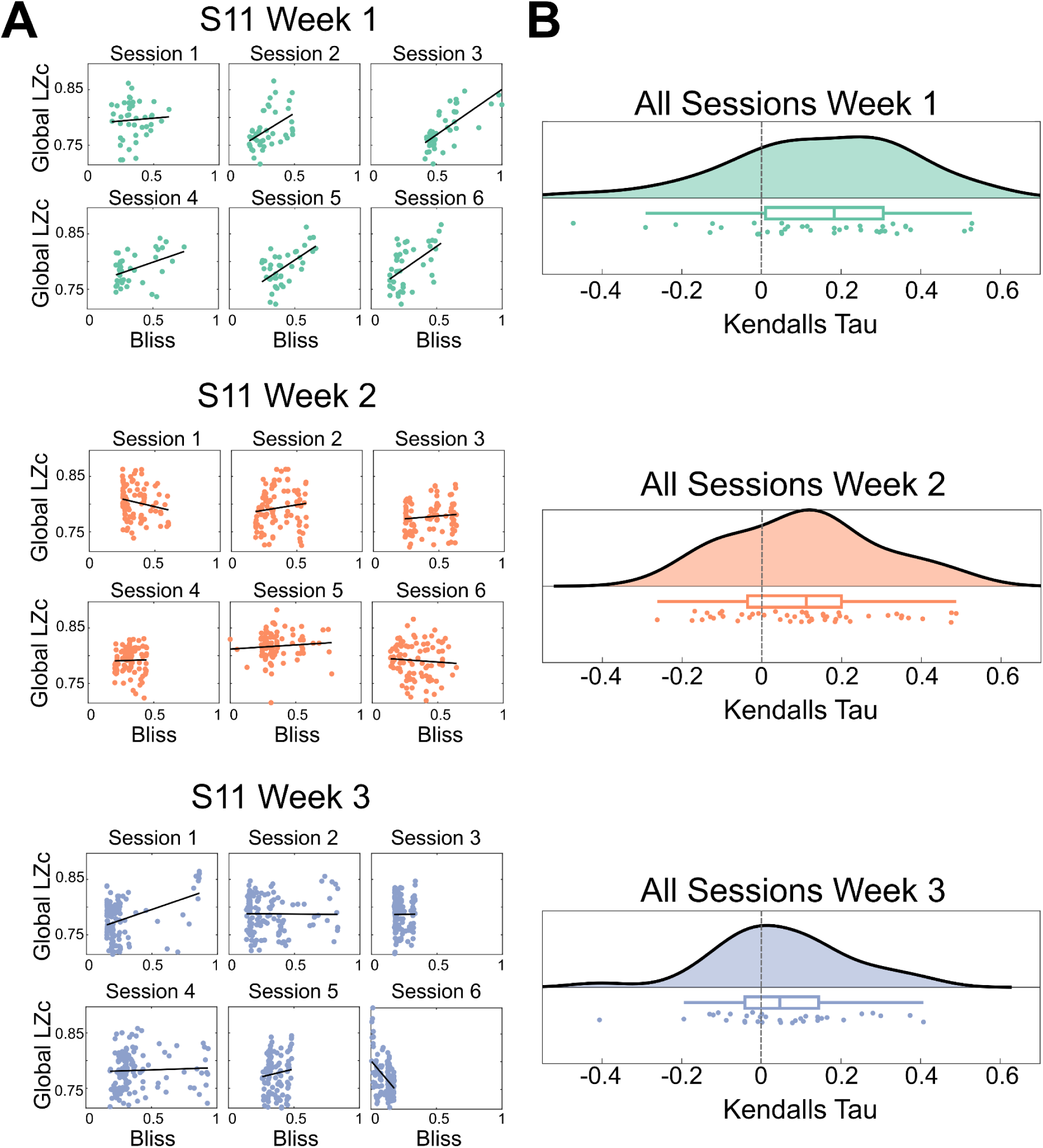
There is a positive relationship between bliss and LZ complexity across weeks. A) Scatterplots of bliss and Global LZc for the first six sessions of weeks 1-3 for subject 11. Lines of best fit were plotted with *lsline* in MATLAB. Plotting the individual sessions highlights the variability between sessions within participants, but exemplifies the enduring positive association between bliss and LZ complexity. B) Raincloud plots highlighting the distribution of every session separated by week. Each data point is the strength of association (Kendall’s tau) between bliss and Global LZc. Linear mixed models found no significant difference between association strength across weeks, with the plots highlighting this general trend towards positive bliss and Global LZc correlations. For the sake of parsimony, only weeks 1-3 are shown.

It should also be noted that the coefficient sizes here are an order of magnitude smaller than some of the data-driven results, suggesting that there is a smaller association between bliss and complexity, when compared to a multivariate neurophenomenological approach. This may arise due to the variance in phenomenology of the sessions; for example, if a participant reported a ‘flat’ intensity of bliss, this would not yield a strong correlation between complexity. This highlights the importance of studying phenomenology from a multivariate perspective, to reveal potentially stronger neurophenomenological effects.

### Stimulus-Driven

As a method to induce an ASC, the phenomenological and neural consequences of breathwork may be contingent on the physiological changes that underpin the practice (Havenith et al., 2024). Previous research has found that blood oxygen saturation decreases as a function of breath retentions, with the lowest blood oxygen saturation at the point in which participants take their first breath (Kox et al., 2014). While the EEG devices were not fitted with a pulse oximeter, accelerometer data was accessible. Therefore, we used data from the z-axis of the accelerometer to identify timepoints in which the participant took their first inhalation after engaging with the breath hold conditions during the breathwork sessions (explained in Methods).

Based on the idea that blood oxygen saturation may be associated with phenomenological changes, we hypothesised that neural data associated with the period in which the participant took their first breath after a breath retention would be significantly different in LZ complexity compared to neural data associated with the period before the start of the breath hold.

### Breath Retentions are Associated with a Small Increase in Lempel-Ziv Complexity

Linear mixed models were used to assess the difference in LZ complexity before and after breath retention. LZ complexity was higher following the breath hold in comparison to the epoch prior to the breath hold commencing in O-O LZ (0.18, SE=0.034, t(1,037)=5.27, p<0.001), F-O LZc (0.16, SE=0.039, t(1,018)=4.06, p=0.002), and Global LZc (0.11, SE=0.034, t(1,011)=3.08, p = 0.037). This tentatively suggests that the breath retentions during the sessions are associated with an increase in neural complexity.

We also hypothesised that the strength of the difference would increase as a function of the intensity of the breathwork session conducted (i.e., peaking at week 3). However, when inputting week as an interaction term, none of the main effects for any spatial complexity type were significant. Further to this, no interaction terms were significant after false discovery rate (FDR) correction.

### Lempel-Ziv Complexity is Modulated by the Amount of Breath Holds

Kox et al. (2014) found that blood oxygen saturation further reduces as a function of the amount of breath holds completed within one session. Therefore, to investigate whether there is a cumulative effect of breath holds on LZ complexity changes, we included the breath hold number as an interaction term. None of the main effects for any spatial complexity type were significant. However, F-O LZc was significant for the interaction terms for the third breath hold (0.32, SE=0.11, t(1,007) = 2.88, p = 0.047) and the fifth breath hold (0.64, SE=0.168, t(1,007) = 3.83, p = 0.003), while the interaction term for the fifth breath hold was significant for O-O LZ (0.42, SE=0.156, t(1,026) = 2.96, p = 0.04; Figure 6). These findings further tentatively suggest that LZ complexity may be associated with the hypoxic conditions associated with breathwork. However, further analyses may be required to more clearly elucidate this relationship.

**Figure 6:**
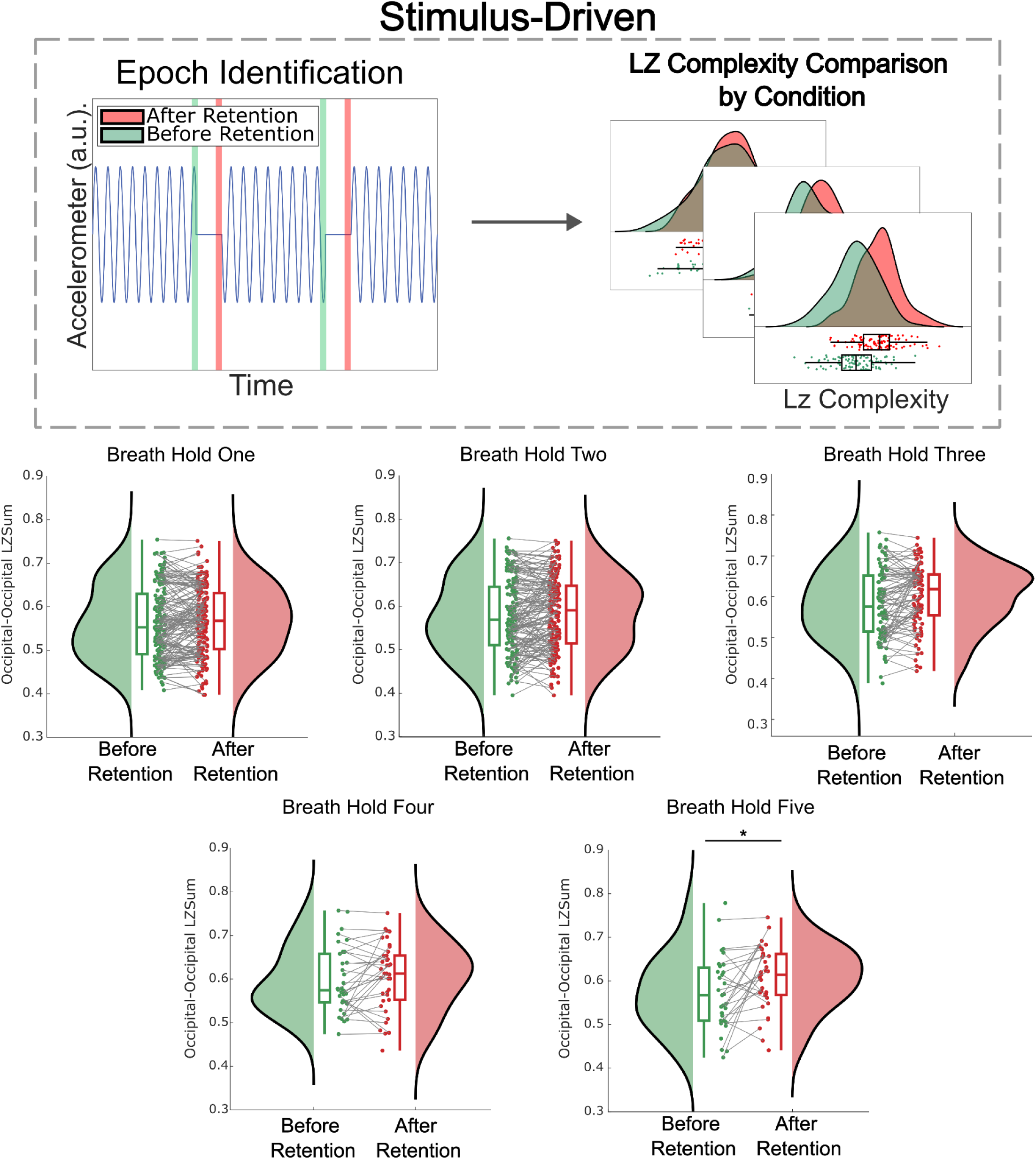
Differences in LZ complexity are heightened due to the amount of breath holds a participant completes during the session. The below raincloud plots are session-level comparisons of occipital-occipital neural complexity both before retention and after retention for each breath hold. As participants increase the number of breath holds, there is a larger difference in neural complexity, with a significant interaction effect for breath hold number five. *p<0.05 FDR corrected.

### Exploratory Analysis: Periodic and Aperiodic Neural Signal Components

As we found significant neurophenomenological associations between psychedelic-like subjective experience states and LZ complexity, we aimed to explore the relationship between phenomenology and other neural signatures associated with psychedelics. Based on the above results demonstrating that there is an increase in LZ complexity in association with bliss and multi-dimensional phenomenological clusters high in subjective experiences closely linked with the psychedelic state, we explored how phenomenology and LZ complexity may relate to aperiodic and periodic components of the EEG PSD.

Based on previous research demonstrating that serotonergic psychedelic administration is associated with a reduction in alpha power (Ville et al., 2016; Pallavicini et al., 2019; Timmermann et al., 2019), we hypothesised that alpha power is negatively associated with the psychedelic-like phenomenological cluster, and the phenomenological dimension of bliss. In addition to this, based on previous research highlighting that LZ complexity outperformed alpha power when correlated with the subjective effects of psilocybin (Mediano et al., 2024), we also hypothesised that the neurophenomenological associations between bliss and LZ complexity will not be explained away by alpha power.

### Alpha Power is Not Robustly Associated with Phenomenological Cluster or Bliss

To explore the potential differences between phenomenological clusters in alpha oscillatory power, a variety of linear mixed models increasing in model complexity were computed (Supplementary Table S2). Most models did not reach (uncorrected) levels of significance at the alpha criterion of 0.05, suggesting that the phenomenological clusters are not robustly associated with alpha power. Further to this, when using the hypothesis-driven approach, most linear mixed models did not find any significant associations between alpha oscillatory power and the phenomenological dimension of bliss (Supplementary Table S3). These results together suggest that there are no robust neurophenomenological associations between alpha oscillatory power and psychedelic-like phenomenology.

### Reduction in Alpha Power and Increase in Global Complexity for Most Breathwork Conditions Compared to Baseline

Due to the lack of robust associations between phenomenology and alpha power, we also explored how alpha power changed as a function of the condition the participant was in. We found that, when compared to the introduction of the session, alpha power was significantly lower in the Breath (–0.106, SE=0.044, t(13,322) = –2.39, p=0.024), Rest (–0.216, SE=0.054, t(13,322) = –3.99, p <0.001), and End (–0.273, SE=0.061, t(13,322) = –4.43, p<0.001), portions of the breathwork sessions. Similarly, when compared to the introduction of the session, Global LZc increased for the Breath (0.15, SE=0.039, t(13,723) = 3.83, p<0.001), Hold (0.228, SE=0.041, t(13,723) = 5.57, p<0.001), Rest (0.286, SE=0.041, t(13,723) = 6.87, p<0.001), and End (0.36, SE=0.048, t(13,723) = 7.41, p<0.001) conditions, and Global LZSum increased for all conditions: Breath (0.32, SE=0.039, t(13,867) = 8.22, p <0.001), Hold (0.434, SE=0.039, t(13,867) = 11.25, p<0.001), Rest (0.39, SE=0.043, t(13,867) = 9.03, p<0.001), Fast (0.335, SE=0.058, t(13,867) = 5.8, p<0.001) and End (0.505, SE=0.049, t(13,867) = 10.41, p<0.001). The data distributions of these are visualised in Supplementary Figure S8. These results concord with similar lines of research demonstrating that alpha power reduces, and LZ complexity increases, as a function of psychedelic administration (Mediano et al., 2024; Schartner et al., 2017; Timmermann et al., 2019; Pallavicini et al., 2019).

### Oscillatory Theta and Beta Power are Modulated in Some Breathwork Conditions

We also explored how the other canonical oscillatory power bands changed as a function of breathwork condition. When compared to the introduction of the session, theta power decreased during the Breath (–0.16, SE=0.044, t(12,149) = –3.49, p<0.001), Rest (–0.21, SE=0.049, t(12,149) = –4.22, p<0.001), and End (–0.25, SE=0.056, t(12,149) = –4.55, p<0.001) conditions, and beta power increased during the Hold (0.11, SE=0.045, t(13,723) = 2.5, p=0.012) condition. Distributions of the data can be found in Supplementary Figure S9.

### Exponent, but Not Alpha, is Associated with Phenomenological Cluster and Bliss

Contextualising these results within a neurophenomenological approach, we explored the potential relationships between (i) Global LZ complexity, (ii) alpha oscillatory power, and (iii) exponent of the PSD as predictor variables, and phenomenological cluster and bliss as response variables. We computed these models to explore whether the exponent of the PSD may also be predictive of both uni-dimensional and multi-dimensional phenomenology. Both Global LZc (0.015, SE=0.006, t(10,298) = 2.70, p = 0.011) and Global LZSum (0.019, SE=0.007, t(10,448) = 2.7, p = 0.009) were significantly associated with bliss. However, only Global LZsum was significantly predictive of the phenomenological cluster (–0.313, SE=0.146, t(12,766) = –2.15, p = 0.045). The aperiodic exponent was significantly associated with bliss (–0.04, SE=0.005, t(10,298) = –7.43, p< 0.001), and phenomenological cluster (0.72, SE=0.116, t(12,607) = 6.25, p<0.001; Figure 7). Alpha power was not significantly associated with phenomenological cluster or bliss.

**Figure 7:**
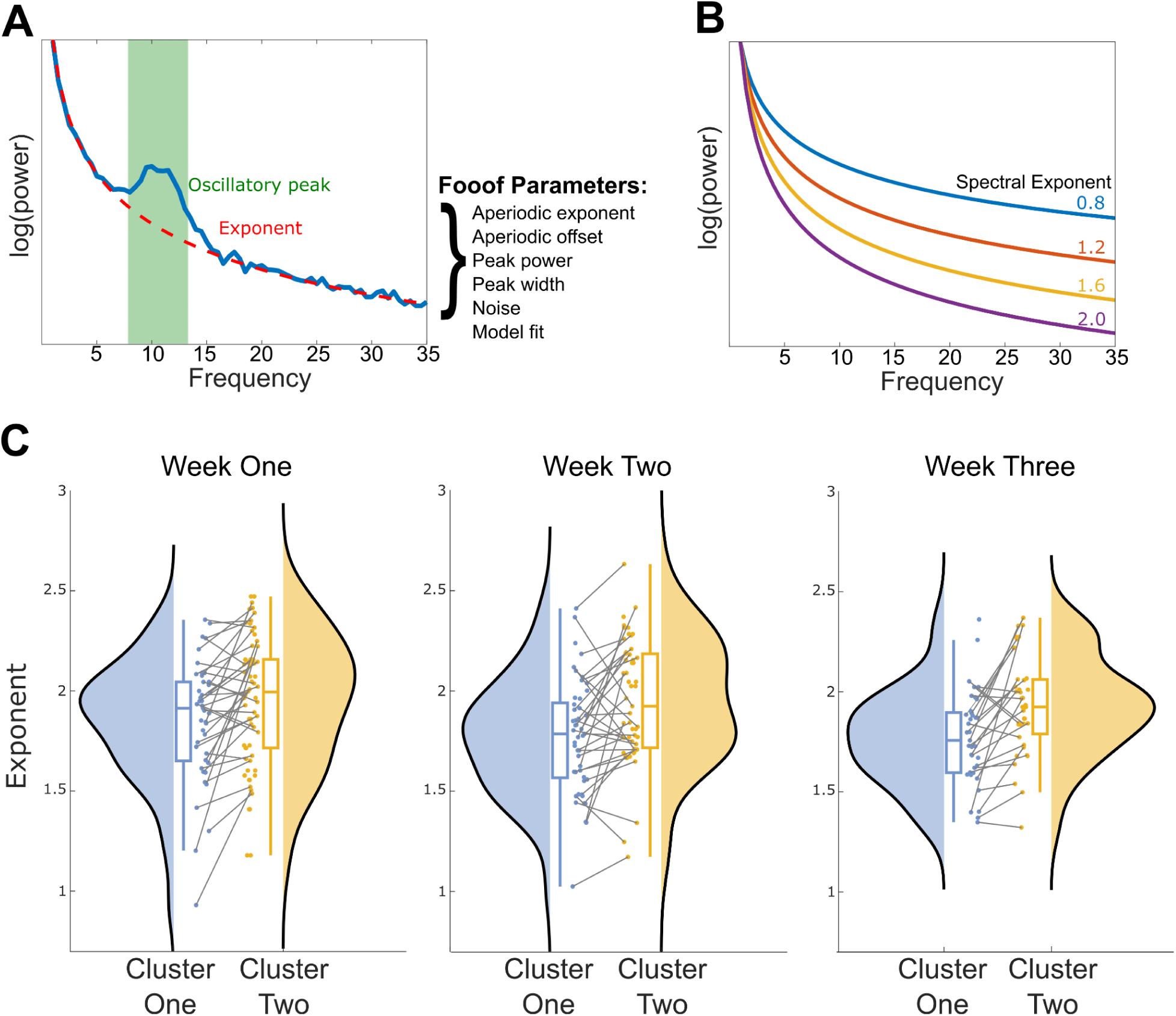
The aperiodic exponent of the PSD shows similar differences between clusters as LZ complexity, where the psychedelic-like cluster (cluster one) is consistently lower in the aperiodic exponent. A) A schematic illustration of the fitting oscillations and one over f (fooof) toolbox. An exemplary neural power spectral density is modelled through the parameterisation of various phenomena, including the aperiodic exponent and the oscillatory peak power. Within this study, we identified neural oscillations as well as parameterised the aperiodic exponent of each neural epoch. B) A schematic example of four separate aperiodic exponent parameters. A higher aperiodic exponent entails a PSD that decays quicker at higher frequencies. C) Session level differences between the average aperiodic exponent for cluster one and cluster two. Cluster one consistently displays a lower aperiodic exponent in comparison to cluster two, even when averaging the data at the session-level.

### No Change in Exponent for Breath Holds

Finally, to explore whether the exponent could successfully distinguish between hypoxic states and non-hypoxic states (as induced by breath holds), we computed linear mixed models in line with the stimulus-driven approach as above, but using the aperiodic exponent as response variables instead of LZ complexity. However, the analysis did not outperform the complexity analyses (as computed above).

## Discussion

In a large-scale neurophenomenological analysis of breathwork as a non-pharmacological ASC, we present several overarching findings. First, we show that phenomenological intensity can be quantitatively captured over time within predetermined experience dimensions using TET. Second, these breathwork-induced experiences separate into two distinct clusters, a positive ‘psychedelic-like’ cluster, and a negative cluster loading onto boredom and physical/mental effort. Third, these distinct phenomenological clusters were also distinct at the neural level, with the psychedelic-like cluster higher in LZ complexity. Finally, while the relationship between LZ complexity and proxy hypoxic state was less robust, we found positive associations between neural complexity and bliss as a psychedelic-like dimension. In exploratory analyses, we found no robust neurophenomenological associations using alpha power, contrary to previous psychedelic research (Timmermann et al., 2019; Mediano et al., 2024). However, the aperiodic exponent of the PSD followed similar neurophenomenological associations as LZ complexity, with flatter PSD distributions in the psychedelic-like subjective experience cluster.

Despite its rich history within Eastern spirituality, the investigation of high-ventilation breathwork practices, where an individual increases their respiratory rate, is in its infancy within contemporary science (Fincham et al., 2023a). Moreover, much like meditation as a broad taxonomic term (Tang et al., 2015; Lutz et al., 2008), breathwork constitutes a plethora of practices (see Fincham et al., 2023a for a review). SOMA breathwork, used within this study, is similar to the Wim Hof method (Kox et al., 2012; 2014), in that it utilises a continuous and standardised breathing pattern in combination with breath retentions to induce a hypoxic state. Both practices have roots in g-tummo meditation (Benson et al., 1982; Benson et al., 1990), a practice from Vajrayana Buddhism combining breathwork and meditation which can induce large core body temperature increases (Benson et al., 1982). These core body temperature increases were strongly associated (r = 0.82) with increases in alpha power in expert practitioners (Kozhevnikov et al., 2013). Within our study, we found converse results; when compared to the introduction of the session as a resting-state baseline, alpha power decreased in the Breath, Rest, and End conditions (Supplementary Figure S8). Further, conscious connected breathwork, a modality comprising an unbroken series of inhalations and exhalations, induced spectral changes in delta, theta, beta 1 (12-15Hz) and beta 2 (15-20Hz) when comparing pre– and post-breathwork resting state data (Bahi et al., 2023). Again, these findings are discrepant from the present study, where we only found within-session decreases in alpha power when comparing the introduction and end of the session (Supplementary Figures S8 and S9). These discrepant results may be due to the potential differences in phenomenological and physiological goals of each practice.

Within the present study, both LZ complexity and the exponent of the PSD were associated with the multidimensional contents of consciousness within breathwork. Unlike the parameterization of oscillatory activity, both LZ complexity and the aperiodic exponent are nonlinear measures of EEG signals, and may measure similar underlying neurophysiological mechanisms; LZ complexity is a quantification of the number of novel patterns within a timeseries, decreasing when the EEG signal is more predictable. LZ complexity changes within ASCs may be related to a change in the aperiodic exponent of the PSD (Colombo et al., 2018), as many states involving a suppression of conscious contents are characterised by a dominance of periodic low frequency activity, thus contributing to a more predictable neural signal (lower in LZ complexity; however see Frohlich et al., 2023 for a discrepant result). Additionally, changes in LZ complexity within the psychedelic state have been ascribed to increases in high-frequency (>25Hz) activity (Mediano et al., 2023), with broadband spectral changes found across a variety of psychedelic substances (Muthukumaraswamy et al., 2013; Carhart-Harris et al., 2016; Pallavicini et al., 2019), thus implicating changes in aperiodic activity (Muthukumaraswamy & Liley, 2018). The overlap between these two measures were also found in linear mixed models (see Supplementary Information), and it is therefore not surprising that they both yielded strong neurophenomenological associations. Relating these findings to the physiology of breathwork, some researchers have theorised that hypoxia and alkalosis induced through ‘high-ventilation’ breathwork practices may cause a shift towards excitability in the excitation/inhibition (E/I) balance (Fincham et al., 2023a), due to the disinhibition of GABAergic inhibitory cortical circuits (Sun et al., 2012; Li et al., 2012). A flattening of the PSD has been ascribed as a marker of a neurophysiological shift towards excitability (Donoghue et al., 2021; Gao et al., 2017), demonstrating that our results support the aforementioned hypothesis with regards to E/I balance. However, other neuroimaging modalities at finer spatial scales (such as ECoG/LFP) are required in order to more fully understand how E/I balance at multiple scales relate to the aperiodic exponent (Ahmad et al., 2022).

It should be noted, however, that LZ complexity is a relatively simplistic computational measure, with information loss due to the binarisation of the EEG timeseries. We used this measure due to the large body of research associating conscious state with LZ complexity (e.g. Aamodt et al., 2021; Casali et al., 2013; Pascovich et al., 2022; Schartner et al., 2015), but highlight that this relationship is less consistent when examining the contents of consciousness (e.g. Aamodt et al., 2023). To further investigate the entropic brain theory’s primary hypothesis, namely that entropy/complexity at the neural level is related to phenomenological richness, future research should examine multiple information theoretical measures within neurophenomenological analyses.

Respiration modulates neural oscillatory activity in animal models (Zelano et al., 2016; Ito et al., 2014; Karalis & Sirota, 2022), is tied to reaction times in sensory-cognitive paradigms (Johannknecht & Kayser, 2022; Nakamura et al., 2018), and is increasingly integrated with theories of emotion, cognition, and conscious experience (Seth, 2013; Seth et al., 2012; Tsakiris & Critchley, 2016). Based on these interdependencies between respiration, cognition, and neural dynamics, we hypothesised that neural complexity would be influenced by the (proxy) hypoxic state. This hypothesis was supported in a subset of LZ complexity types, where there was a small increase in complexity after breath retention compared to before. Further, we found that the number of breath holds conducted within a single session had a cumulative impact on LZ complexity (Figure 6). These results suggest a relationship between the physiological state of the participant and neural complexity, just as recent research has found a relationship between the experiential depth within multiple breathwork modalities and CO_2_ saturation (Havenith et al., 2024). However, it should be stated that the method used to identify epochs associated with the release of the breath within this research is exploratory. We may not be certain that the most abrupt changes in accelerometer signal are not simply movement artifacts. Future research may be necessary to validate the accuracy of this methodology.

While we remain in relatively seminal research stages, evidence is increasingly highlighting the transdiagnostic clinical efficacy of psychedelic-assisted psychotherapy (PAP; Goodwin et al., 2022; Carhart-Harris et al., 2021; Bogenschutz et al., 2022; Gukasyan et al., 2022), with psychedelic phenomenology a strong predictor of clinical outcomes (Yaden & Griffiths, 2020; Garcia-Romeu et al., 2015; Roseman et al., 2018; Ko et al., 2022; however, see Olson, 2020). Additionally, (slow-paced) breathwork interventions may be associated with moderate-to-small differences in stress, depression, and anxiety when compared to controls (in non-clinical populations; Fincham et al., 2023b). Due to the potential neural and phenomenological similarities between breathwork and the psychedelic experience, we advise researchers to investigate breathwork as a non-pharmacological, psychedelic-adjacent clinical intervention, with care and investigative attention given towards phenomenology, set and setting (Hartogsohn, 2017; Carhart-Harris et al., 2018), and embedding the experience(s) within a psychotherapeutic paradigm (Gründer et al., 2023). We also highlight that, if breathwork is a scalable intervention, it is also imperative to investigate the proclivity of adverse events when implemented clinically. TET can also track long-term symptom dynamics (Niedernhuber et al., 2023), moving away from discrete sampling and towards a more comprehensive approach to the study of mental health disorders (Lewis-Healey et al., 2022).

While this study leverages a powerful neurophenomenological within-participant design, some limitations should be mentioned. First, the lack of control group prevents us from ascribing respiratory modulation to phenomenological changes within breathwork. While it was not deemed essential to include a control group within this neurophenomenological study, it will undoubtedly be necessary for future clinical trials investigating breathwork, and ample thought is required to determine what sufficient controls and blinding would look like (Nayak et al., 2023; Muthukumaraswamy et al., 2021; Van Dam et al., 2018). Additionally, while we have collected a highly powered dataset, we recommend that future research should attempt to investigate breathwork at the neurophenomenological level with larger sample sizes, so we may gain a deeper insight into the generalisability of these findings. Further studies should be conducted in laboratory settings to measure the specific oxygen exchange, breath volumetry, and general vital signs in order to physiologically characterise this now standardised technique. Second, at the inception of this study, there was little phenomenological research investigating breathwork as an ASC. There may, therefore, be phenomenological elements central to the practice that have been missed, such as the intensity of felt bodily sensations. We thus present our phenomenological dimensions as foundational work into this breathwork modality, and reiterate the necessity of the collection of qualitative data (Lawrence et al., 2022). A third limitation regards the low-density EEG headsets used; a lack of spatial information here prevents us from drawing further similarities between breathwork and psychedelics at the neural level. For example, with a limited number of channels in the EEG headset, more comprehensive functional connectivity analyses would not be applicable to this dataset, as have been applied to psychedelics (Barnett et al., 2020; Tagliazucchi et al., 2016; Carhart-Harris et al., 2017). In addition to this, we also acknowledge that the portable EEG headsets used promoted a more conservative preprocessing pipeline. An ICA was more difficult to apply to this dataset due to the low-density nature of the headsets, which may have therefore influenced the prevalence of movement artifacts within the dataset. We therefore highlight the importance of replicating these findings with high-density EEG systems in more controlled laboratory conditions to mitigate the potential effects of movement artifacts. Finally, it should be noted that there may be other explanations with regards to the interpretation of the phenomenological clusters. For example, due to the relatively higher dimensions of Meta-Awareness, Clarity, and lower dimensions of Physical and Mental Effort in Cluster One, this could be interpreted as relating to mindful qualities, as all of these dimensions are found in the phenomenological matrix of mindfulness (Lutz et al., 2015). This could also fit into previous research demonstrating that meditation may increase the entropy of neural dynamics (Vivot et al., 2020)^4^. Again, future qualitative research is necessary to uncover the phenomenological nuances between breathwork, psychedelics, and meditation, which may provide a more accurate decomposition of phenomenological clusters in future research.

In summary, our study demonstrates that breathwork may yield neural and phenomenological consequences similar to serotonergic psychedelics. Both LZ complexity and the exponent of the neural PSD were associated with psychedelic-like subjective experience states, whereas alpha oscillatory power did not yield robust neurophenomenological associations. While further research is required to gain a more comprehensive mechanistic understanding of breathwork, our findings and experimental framework form a path to simultaneously evaluate neural and phenomenological dynamics. Placing both phenomenological (TET) and neural data in the same analytical quantitative space has “narrow[ed] the epistemological and methodological distance in cognitive neuroscience between subjective experience and brain processes” (Lutz & Thompson, 2003, p.49), one of the ultimate goals of neurophenomenology. We encourage researchers to further explore the effects of breathwork at the subjective and neural level, and how it may be implemented as a non-pharmacological ASC within clinical paradigms.

## Acknowledgements

We wish to thank Vanessa Potter for help in shaping the experiential dimensions of this project. We also wish to thank Niraj Naik for creating the breathwork course used within this study, and for providing online instruction and guidance for participants’ throughout the study. This work was supported by Human Experience Dynamics consultancy funds (PJJR.GAAA) to Tristan Bekinschtein and an ESRC DTP fellowship to Evan Lewis-Healey.

## Footnotes

1 It is of note that this association may be most robust within high temporal resolution neuroimaging modalities, such as EEG and MEG. A recent study has found conflicting results when investigating the relationship between neural complexity and psychedelics using fMRI (McCulloch et al., 2023).

2 Unlike certain meditative states, which may require tens of thousands of hours of practice (Yang et al., 2024).

3 We acknowledge that separate peaks in the alpha frequency band may represent separate underlying neural mechanisms (Cohen, 2017). However, to simplify the interpretation of the results for the output of this paper, we focus purely on the relative power of the alpha frequency band through the summation of multiple oscillatory peaks if and when they occurred.

4 However, the relationship between neural entropy/complexity and the meditative state is nuanced, complex, and contingent on the meditative style practised (Atad et al., 2023).

## Notes

### Competing Interest Statement

The authors have declared no competing interest.

### Summary of Updates

Parts of introduction have been updated to clarify TET; Figure 4 updated; Parts of discussion extended to clarify on future directions and limitations of study; further analysis conducted and reported on in data-driven section; overall paper shortened

