## Supplementary Information for "Breathwork-Induced Psychedelic Experiences Modulate Neural Dynamics"

**Alpha Power and Exponent are Associated with Global LZ Complexity**

To explore the relationship between alpha power, Global LZ complexity, and the aperiodic exponent, we computed linear mixed models between these variables. First, we computed a linear mixed model to examine the association between Global LZ complexity and alpha oscillatory power. We found that alpha was positively associated with Global LZc (0.077, SE=0.018, $t(13{,}075) = 4.35$, $p<0.001$) and Global LZsum (0.123, SE=0.017, $t(13{,}205) = 7.40$, $p<0.001$). To examine whether this association survived after adjusting for the aperiodic exponent, we input (i) alpha oscillatory power, (ii) the exponent of the neural PSD, and (iii) week, in the same linear mixed model, with Subject, Week and Session as nested random effects. The association between alpha power and complexity reduced, but remained significantly associated for both Global LZc (0.044, SE=0.017, $t(12{,}608) = 2.61$, $p = 0.01$), and Global LZSum (0.05, SE=0.009, $t(12{,}767) = 5.52$, $p <0.001$). Further to this, the exponent was significantly negatively associated with Global LZc (-0.25, SE=0.014, $t(12{,}608) = -16.75$, $p < 0.001$) and Global LZsum (-0.53, SE=0.023, $t(12{,}767) = -22.65$, $p < 0.001$), where the slope of the PSD decreases as LZ complexity increases. This analysis is included in the supplementary material as it simply highlights how LZ complexity and the aperiodic exponent have significant overlap, due to the mathematical consequences with regards to how both measures are computed.

Fig S1: An example temporal experience trace (TET) graph from week three of the breathwork course, where a subject retrospectively traced the subjective intensity (solid black line) of bliss. Dotted vertical lines along the x-axis served as cues regarding instructions of the breathwork session, where S=Start [breathing], and H = Hold [breath].

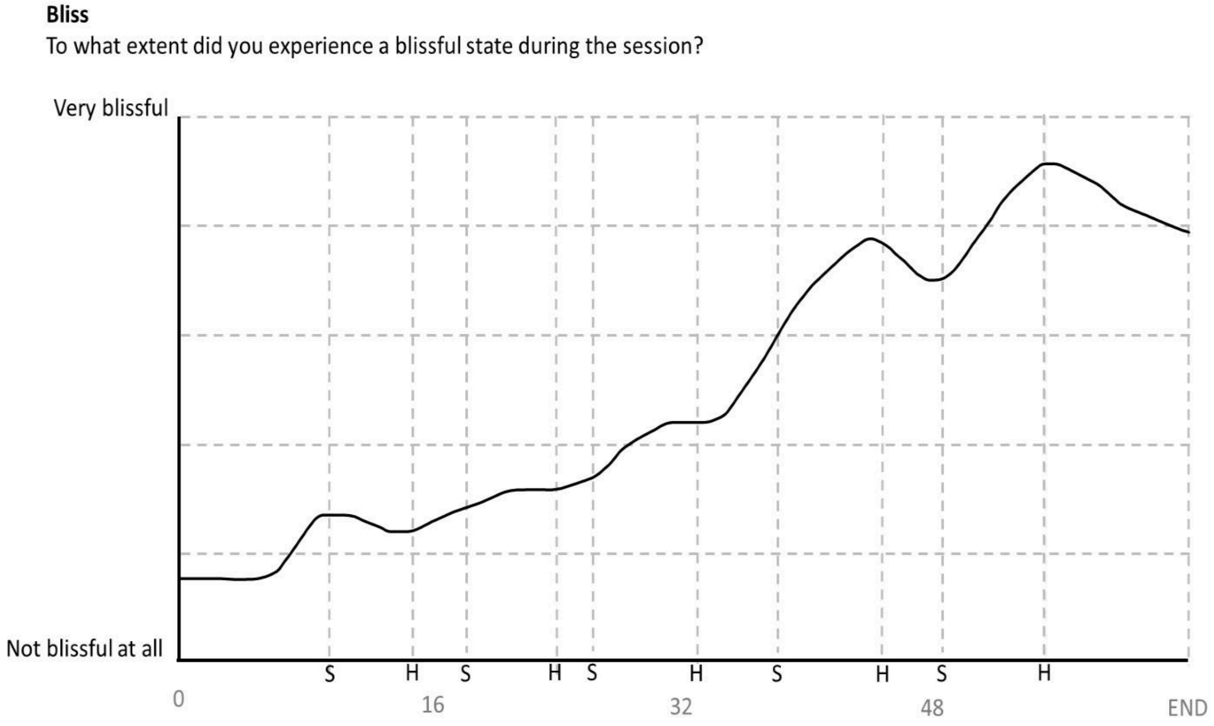

Fig S2: Total number of sessions contributed to the TET dataset. These sessions were used for the k-means clustering of the TET data (301 sessions).

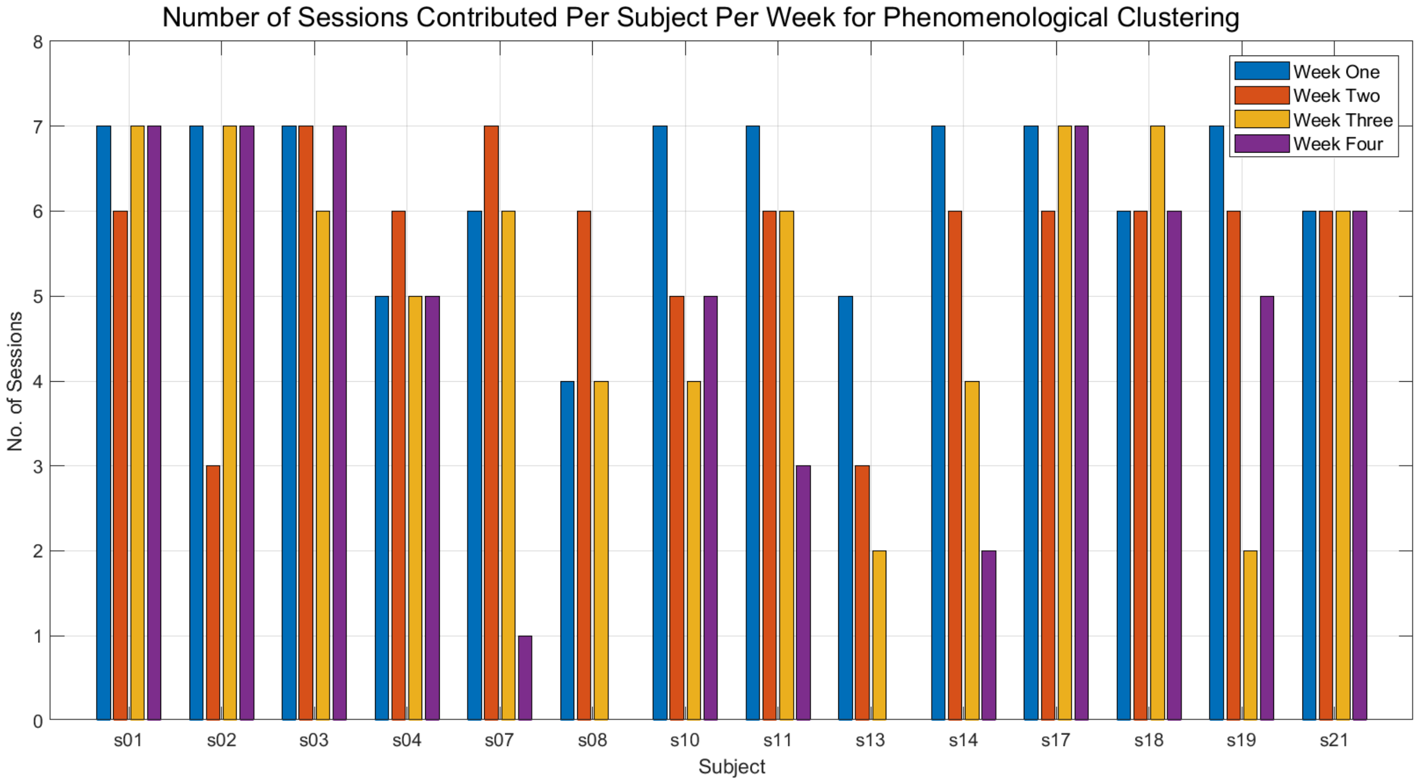

Fig S3: Number of sessions contributed to the data-driven neurophenomenological analyses. The sessions below represent the number of EEG sessions contributed to the analysis per participant per week, where over 20% of the neural data remained after cleaning.

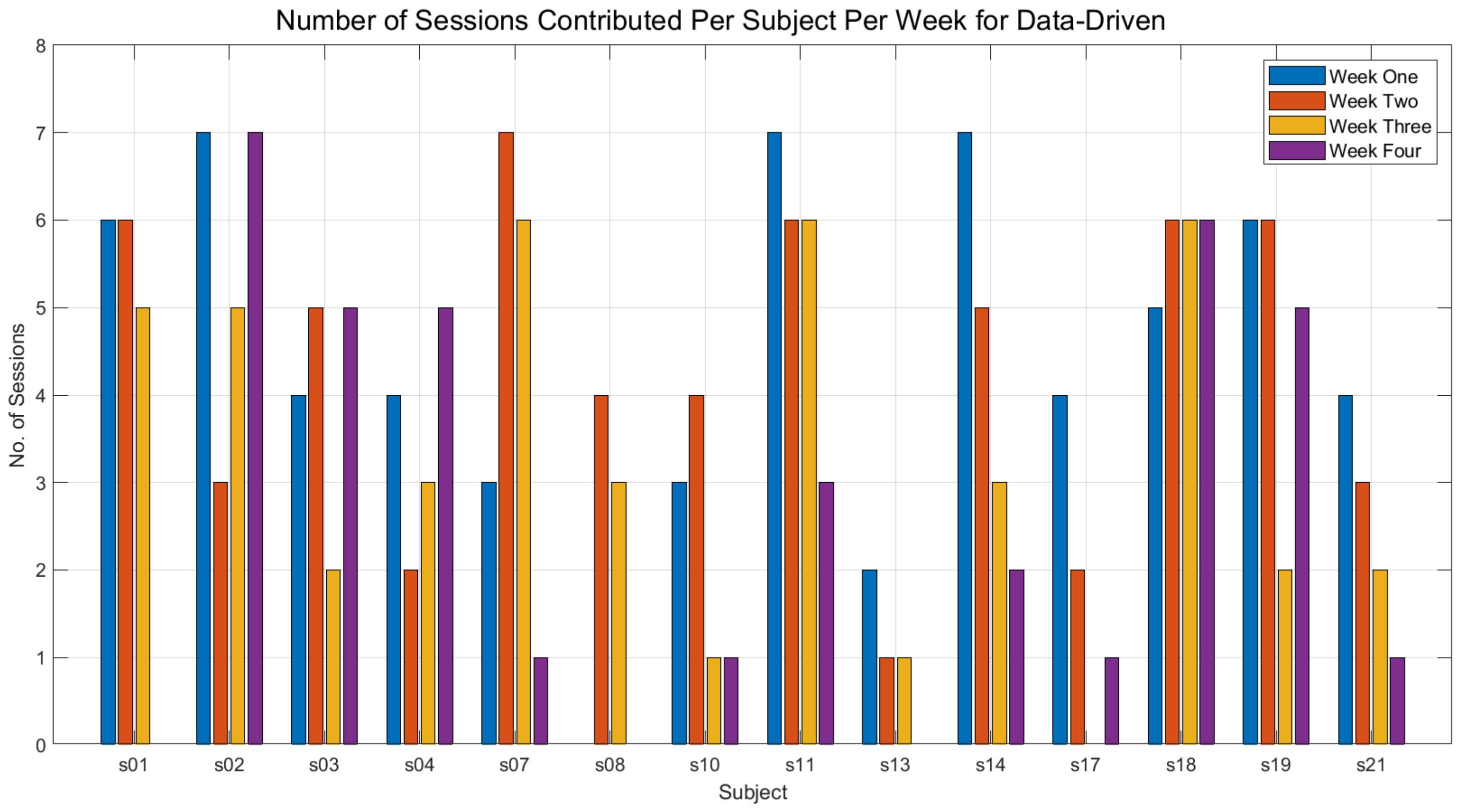

Fig S4: Number of EEG sessions contributed to the hypothesis-driven neurophenomenological analysis. These represent neural data with more than 50% of the epochs remaining after cleaning.

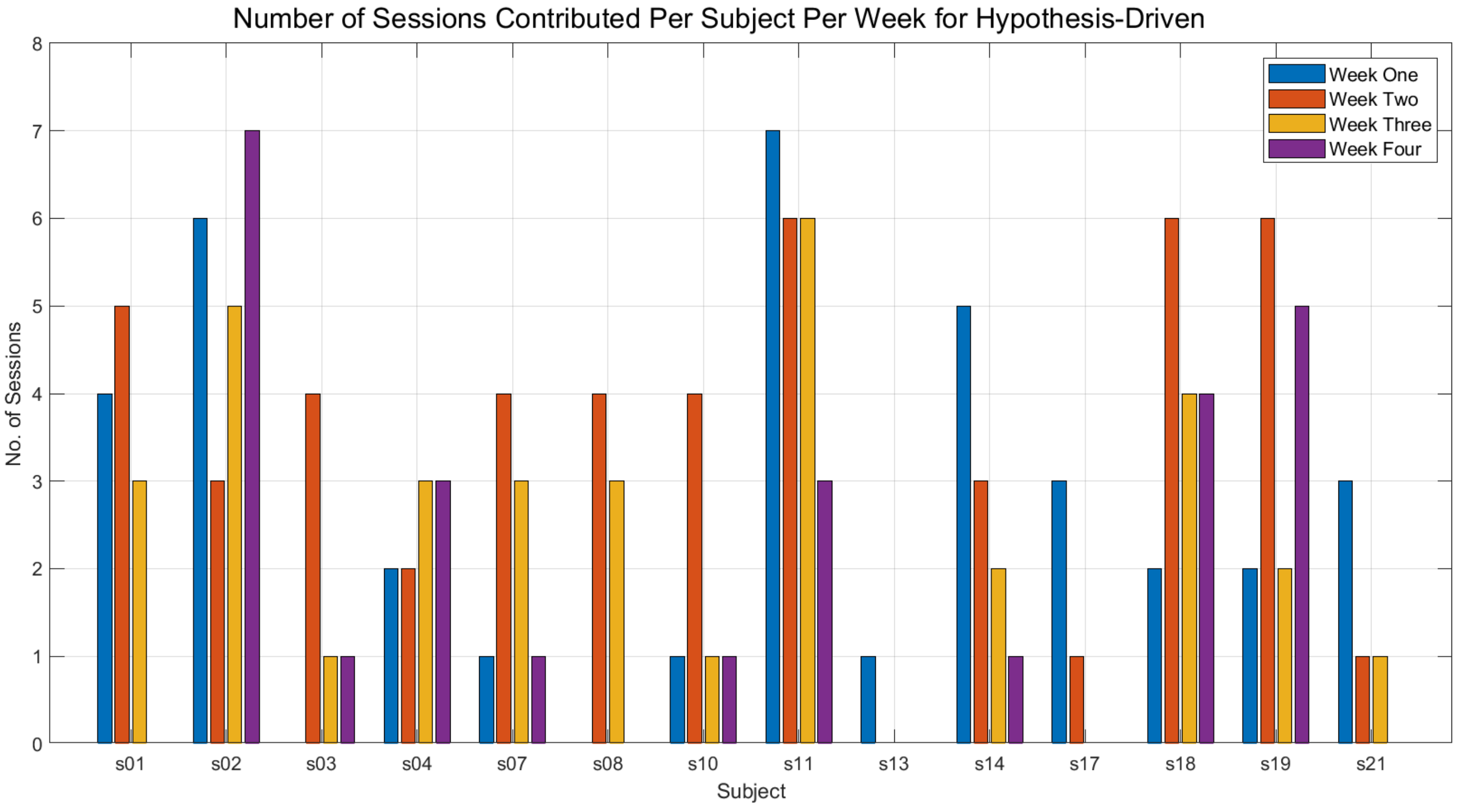

Fig S5: Number of EEG sessions contributed to the stimulus-driven analyses, investigating the relationship between physiological state and neural complexity.

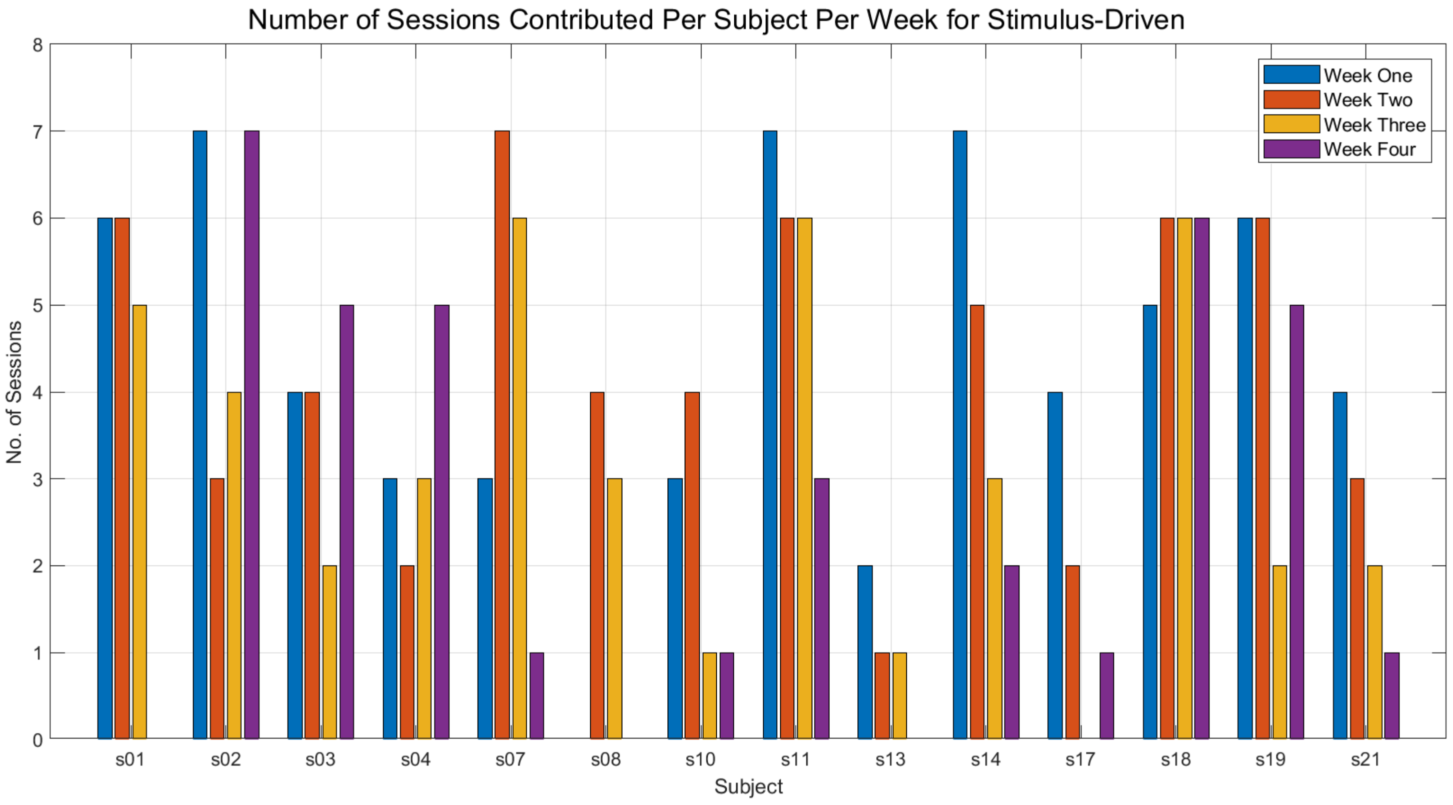

Figure S6: Session-level comparisons between cluster one and cluster two for week four of the breathwork course. On the left are the Global LZSum values, and on the right are the aperiodic exponent values. Each datapoint represents the average value for that session, with lines connecting opposing clusters that occurred within the same session. Within week four, there were more single cluster sessions than previous weeks.

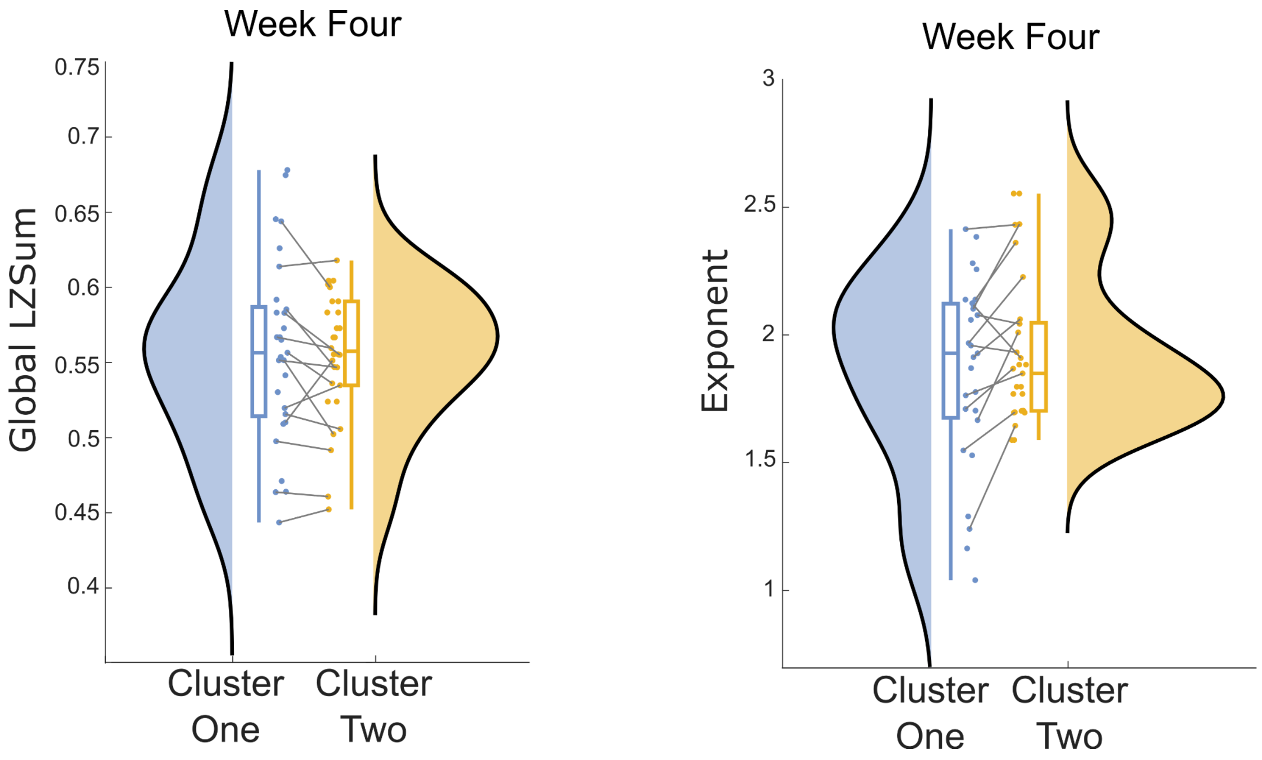

Figure S7: The main and interaction effects for the linear mixed models computed within the hypothesis-driven approach. *p<0.05; **p<0.01; ***p<0.001.

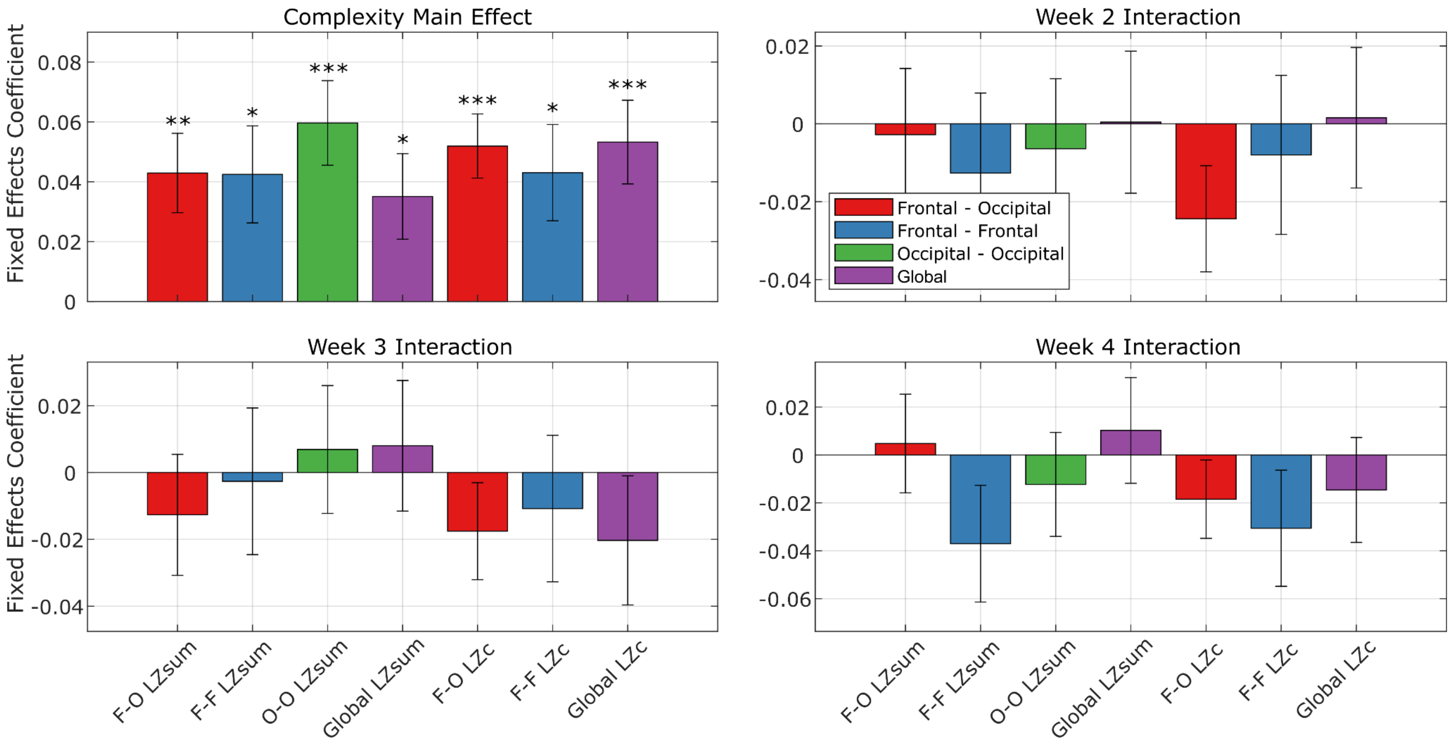

Figure S8: The distribution of Global LZSum, Bliss, aperiodic exponent, and alpha power by condition. Each panel is a corresponding neural or phenomenological feature from the whole dataset. When computing linear mixed models to compare the difference in conditions, every condition was significantly different to the introduction of the sessions at p<0.001 (FDR uncorrected) for global LZSum, bliss, and the aperiodic exponent. For alpha, only the breath (p=0.017), rest (p<0.001), and end (p<0.001) conditions were significantly different from the introduction.

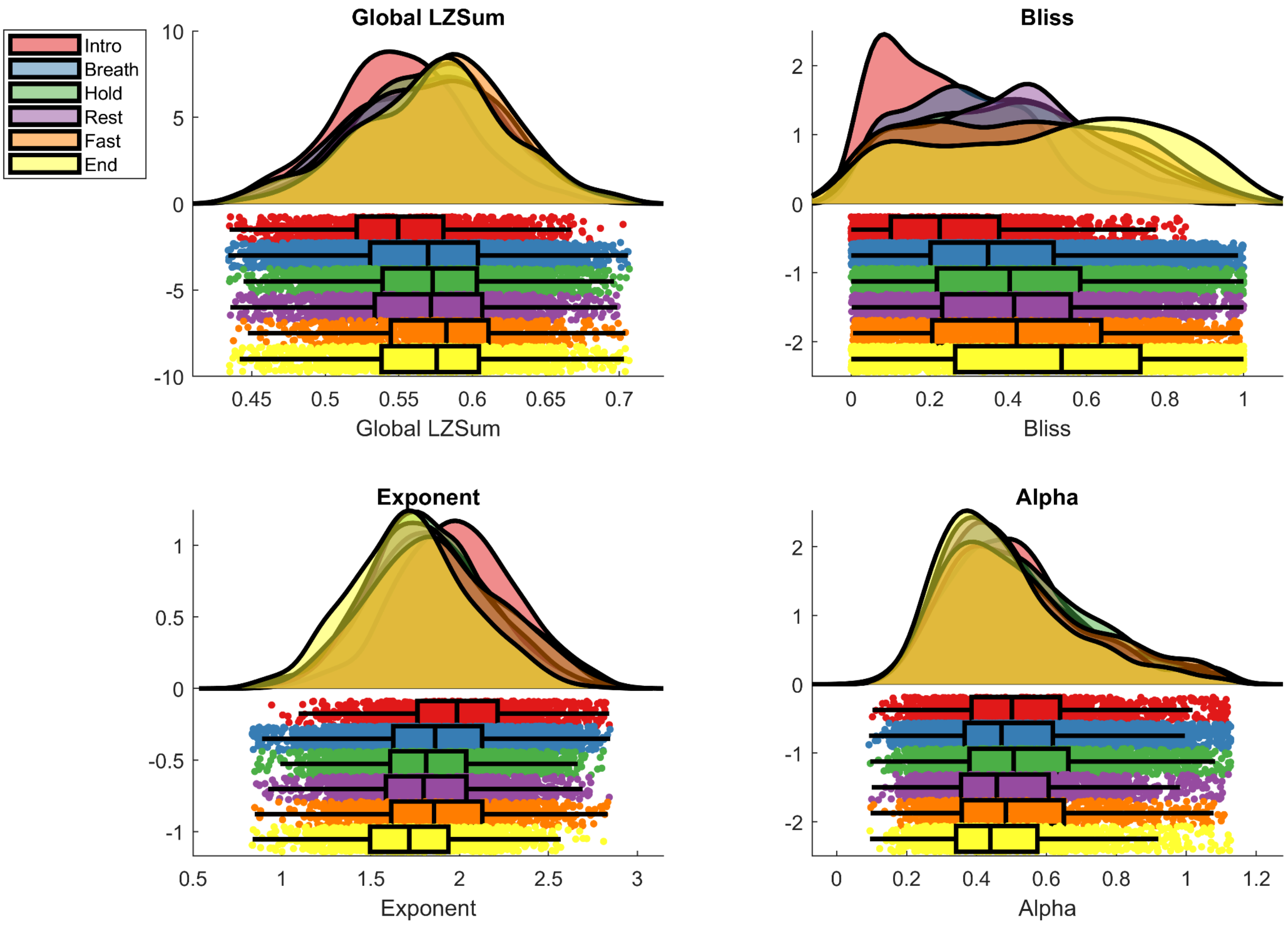

Figure S9: The distribution of delta, theta, and beta oscillatory power by condition, as in Figure S8. Linear mixed models were used to statistically analyse the difference between the introduction of the session and each corresponding cognition of the breathwork session. Significant differences were only found between the Introduction and Breath, Rest, and End conditions for theta (all p<0.001), and the introduction and hold conditions for beta (p=0.012). *p<0.05; ***p<0.001.

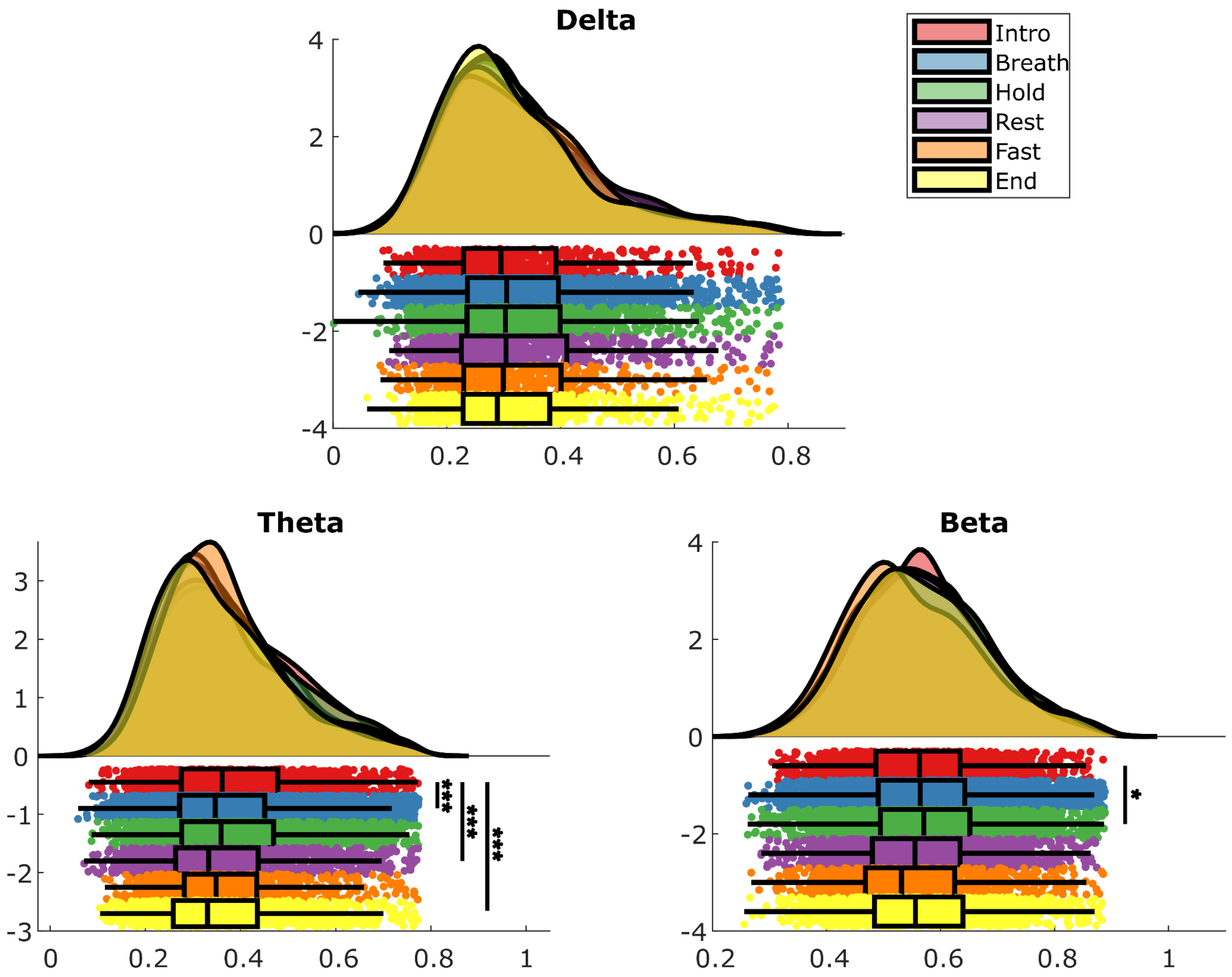

Figure S10: A comparison of the distribution of Global LZSum values per subject by cluster. This figure serves to illustrate the differences by subject in terms of neural complexity and phenomenology.

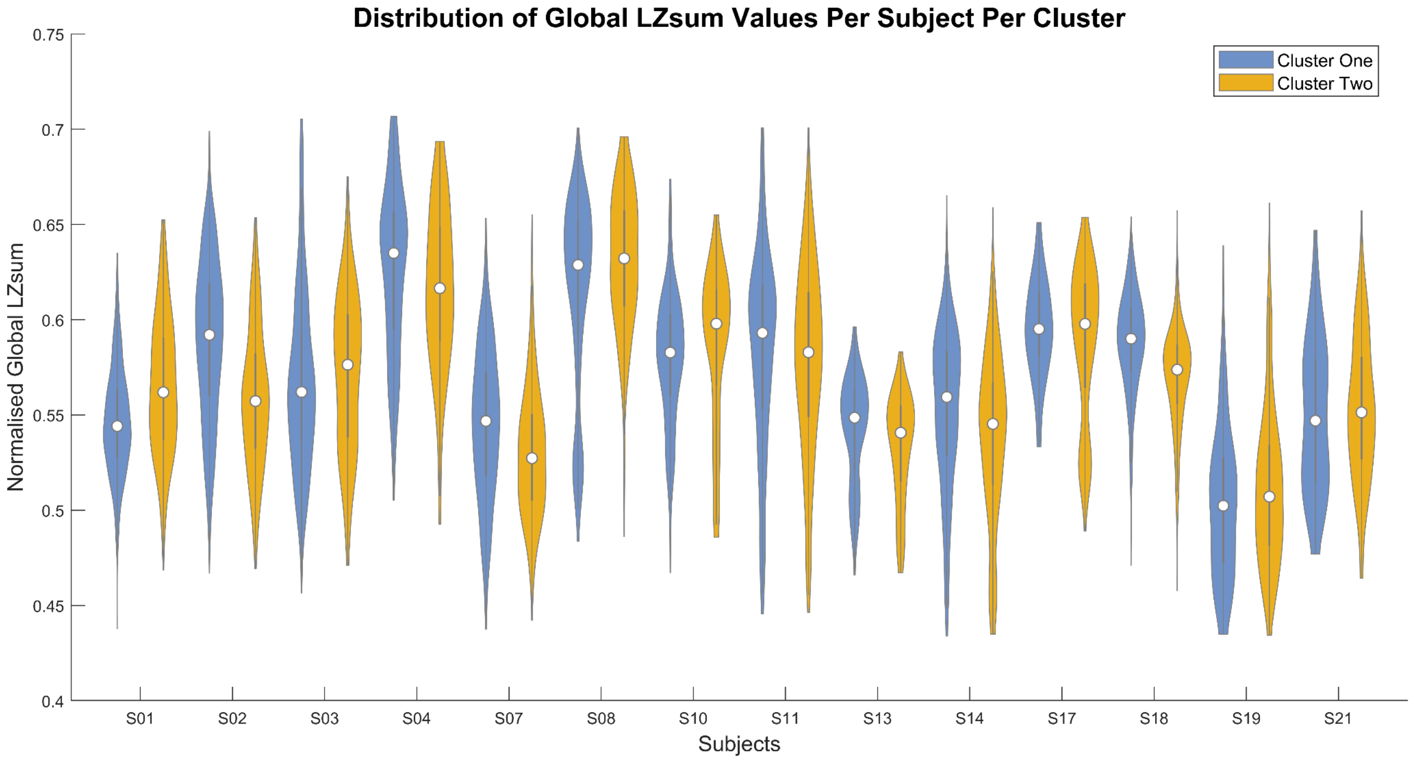

Figure S11: A comparison of the within-session log odds ratio of cluster one prevalence by week. This figure highlights the distribution of odds ratio values by session for each week, where positive values indicate a higher reporting of Cluster 1 towards the end of the session compared to the beginning. There were no significant differences between weeks when comparing the log odds ratios on a within-session basis.

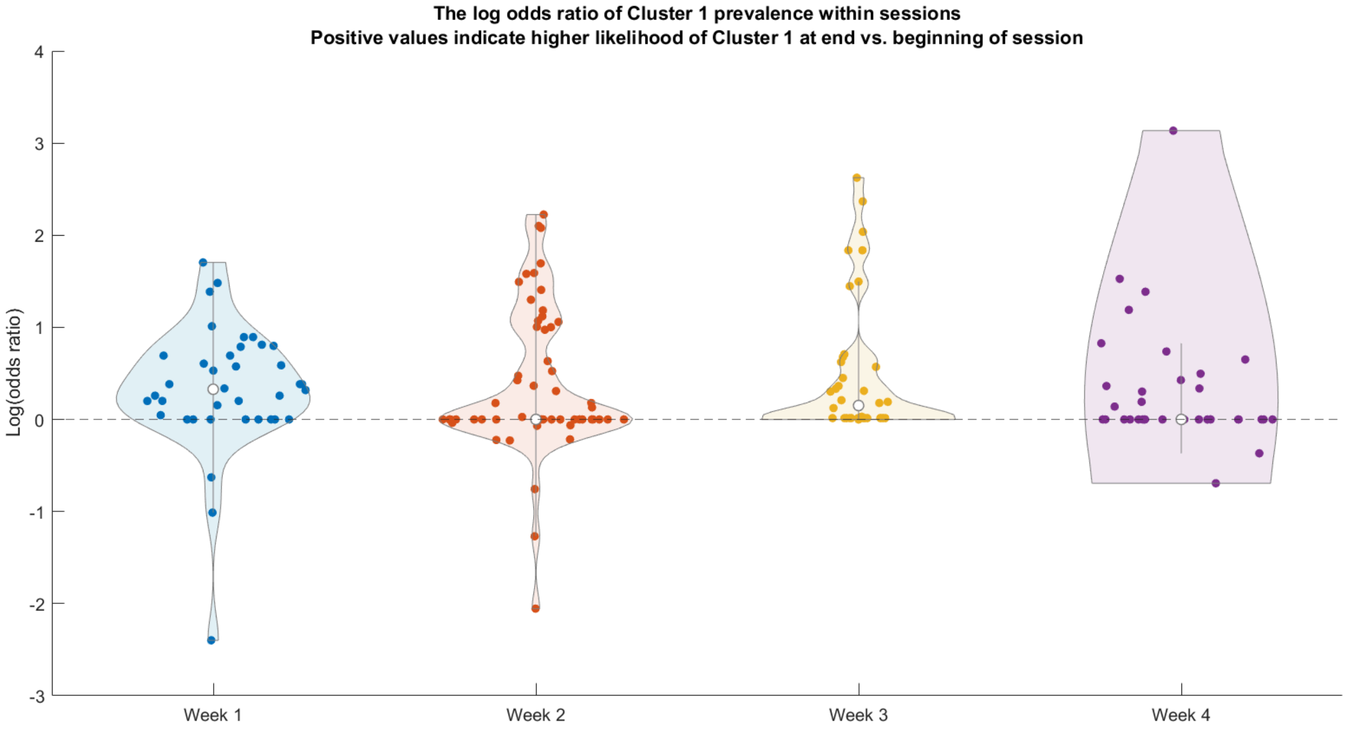

Figure S12: A comparison of the proportion of time spent in each cluster by subject. Using the data within this figure, a paired sample t-test found no significant difference between the proportion of time spent in each cluster (p = 0.47).

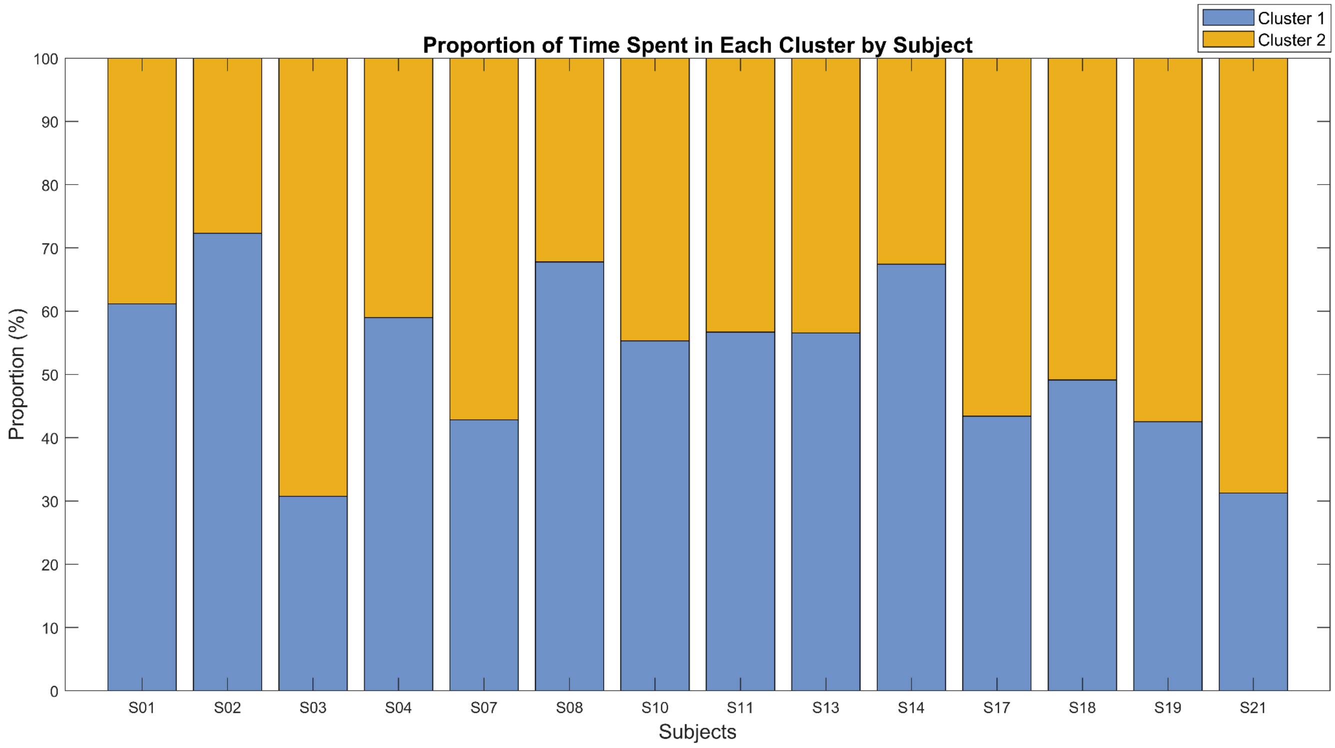

Figure S13: A comparison of the session-level intercepts and slopes for each subject within a sample data-driven model with the formula 'global_lzsum ~ Cluster + (Cluster | Subject:Week:Session)'. The ICC for this statistical model is 0.54 (Table S4), indicating high variance between groups. This solidifies the necessity for factoring in the nested structure of the experimental design within the statistical model (Musca et al., 2011).

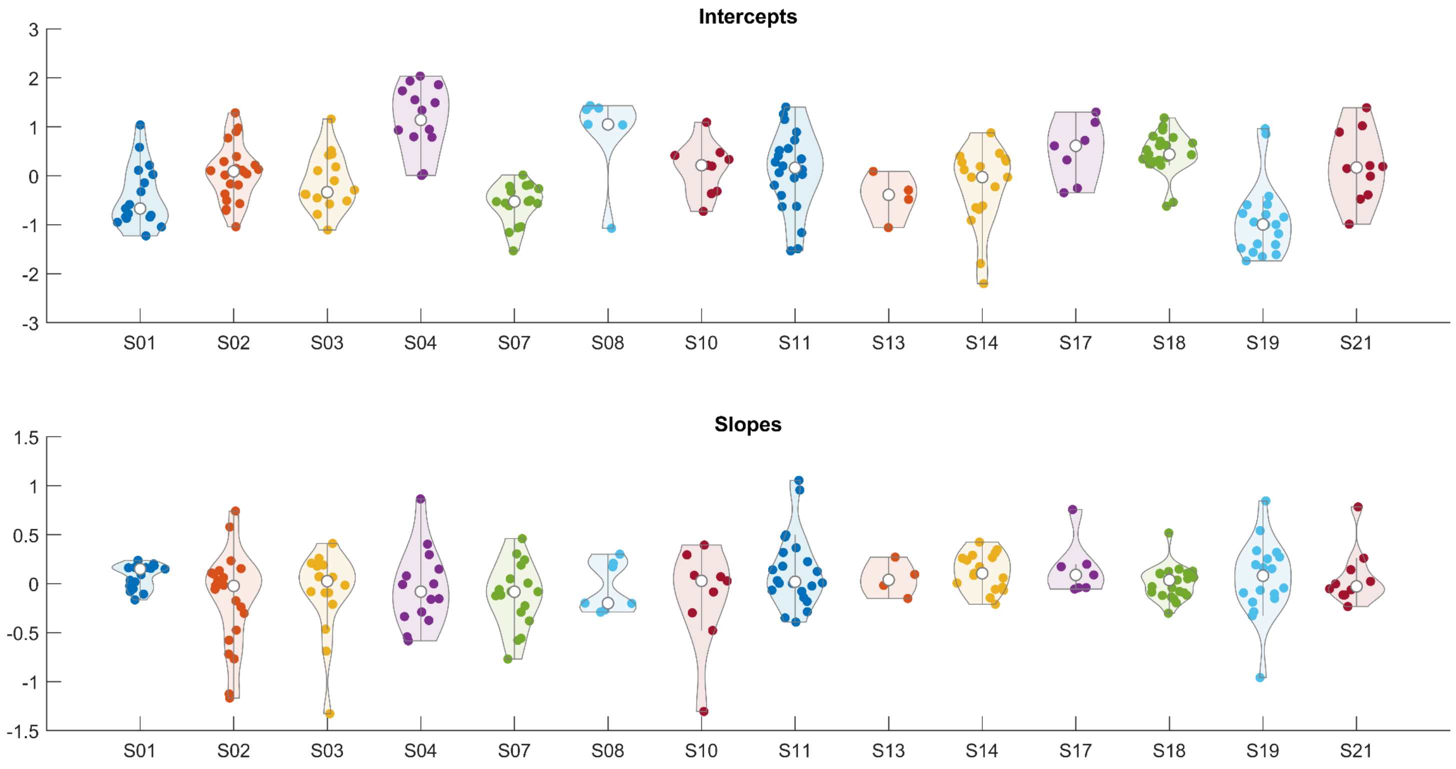

Figure S14: The estimated intercepts (top) and slopes (bottom) for the exponent from the generalised linear mixed model using a logit link function with the following formula: 'Cluster ~ 1 + Week + global_lzsum + global_lzc + alpha + exponent + (1 + global_lzsum + global_lzc + alpha + exponent | Subject:Week:Session)'. The ICC of this model was 0.44.

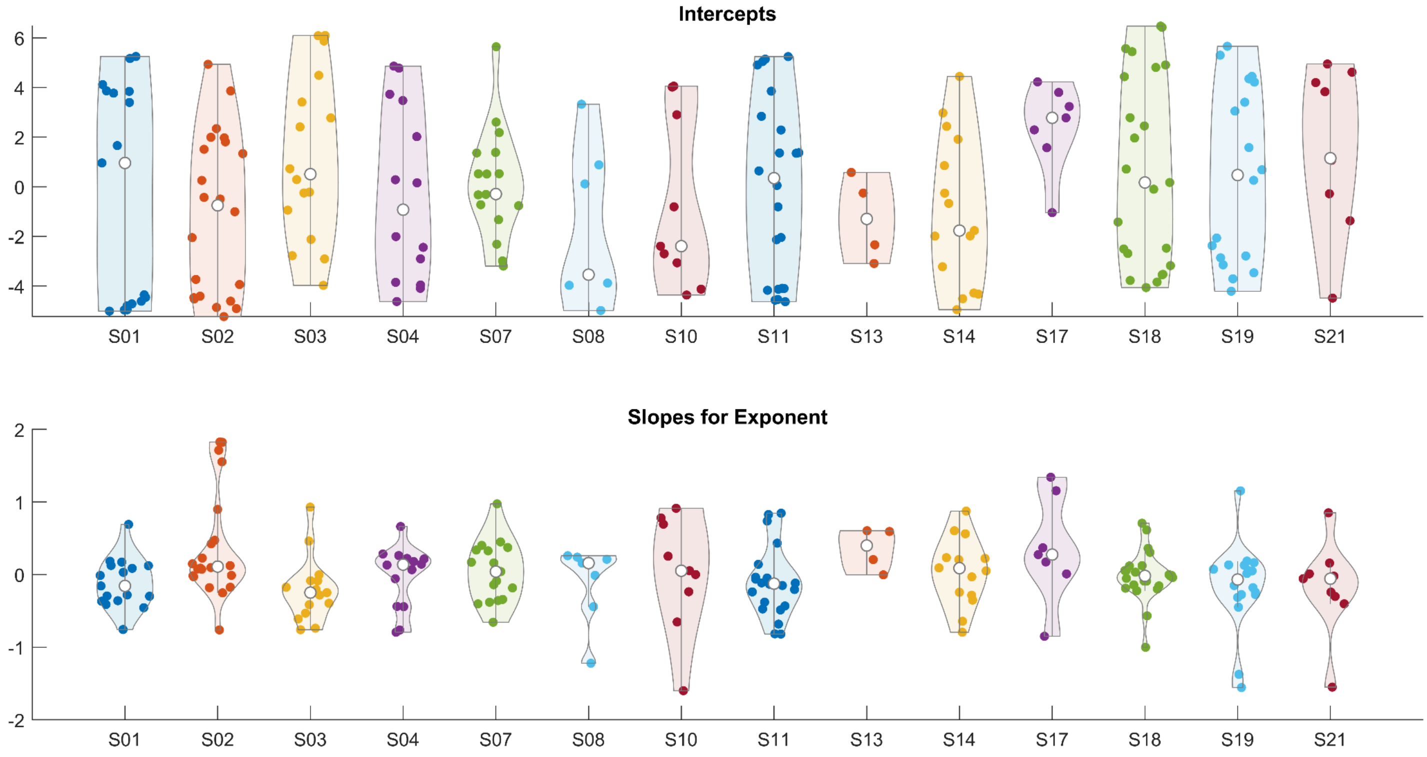

Table S1: Differences in Intensities of Clusters by Phenomenological Dimension. Each row is a value of an output from a linear mixed model, with the overarching formula 'Predictor ~ Cluster + (Cluster | Subject:Week:Session)'. Negative coefficients represent lower intensity values in Cluster Two for the corresponding dimension.  SE: Standard Error; BIC: Bayesian Information Criterion.

| **Dimension** | **Coefficient** | **SE** | **P-value** | **BIC** |
|---|---|---|---|---|
| Meta Awareness | -0.06 | 0.015 | <0.001 | -35364 |
| Presence | -0.25 | 0.011 | <0.001 | -34290 |
| Physical Effort | 0.07 | 0.013 | <0.001 | -43536 |
| Mental Effort | 0.15 | 0.012 | <0.001 | -37812 |
| Boredom | 0.17 | 0.013 | <0.001 | -43799 |
| Receptivity | -0.2 | 0.012 | <0.001 | -45823 |
| Emotional Intensity | -0.14 | 0.012 | <0.001 | -42661 |
| Clarity | -0.18 | 0.012 | <0.001 | -42664 |
| Release | -0.21 | 0.012 | <0.001 | -38509 |
| Bliss | -0.24 | 0.01 | <0.001 | -35013 |
| Embodiment | 0.01 | 0.015 | 0.50 | -38693 |
| Insightfulness | -0.16 | 0.01 | <0.001 | -44688 |
| Anxiety | 0.09 | 0.001 | <0.001 | -52667 |
| Spiritual Experience | -0.2 | 0.01 | <0.001 | -40455 |

Table S2: Linear mixed models examining the difference in alpha oscillatory power between clusters. Coefficient values and associated metrics are reported, with the formula increasing in model complexity. P-values are uncorrected. SE: Standard Error; BIC: Bayesian Information Criterion.

| **Formula** | **Coefficient** | **SE** | **P-value** | **BIC** |
|---|---|---|---|---|
| alpha ~ 1 + Cluster + (1 \| Subject) | -0.014 | 0.003 | <0.001 | 32841 |
| alpha ~ 1 + Week + Cluster + (1 \| Subject) | -0.008 | 0.003 | 0.008 | 32601 |
| alpha ~ 1 + Cluster + (1 + Cluster \| Subject) | -0.016 | 0.016 | 0.30 | 32489 |
| alpha ~ 1 + Week + Cluster + (1 + Cluster \| Subject) | -0.011 | 0.015 | 0.44 | 32317 |
| alpha ~ 1 + Week + Cluster + (1 + Cluster \| Subject:Week) | -0.006 | 0.013 | 0.63 | 31345 |
| alpha ~ 1 + Cluster + (1 \| Subject:Week:Session) | 0.01 | 0.004 | 0.009 | 28673 |
| alpha ~ 1 + Week + Cluster + (1 + Cluster \| Subject:Week:Session) | 0.009 | 0.009 | 0.30 | 28443 |
| alpha ~ 1 + Cluster + (1 + Cluster \| Subject:Week:Session) | 0.008 | 0.009 | 0.35 | 28423 |
| alpha ~ 1 + Week*Cluster + (1 + Cluster \| Subject:Week:Session) | 0.002 | 0.016 | 0.91 | 28470 |
| alpha ~ 1 + Week*Cluster + (1 \| Subject:Week:Session) | 0.011 | 0.009 | 0.21 | 28709 |
| alpha ~ 1 + Condition + Cluster + (1 + Cluster \| Subject:Week:Session) | -0.002 | 0.009 | 0.83 | 28244 |
| alpha ~ 1 + Condition*Cluster + (1 + Cluster \| Subject:Week:Session) | 0.041 | 0.011 | <0.001 | 28252 |

Table S3: Linear mixed models examining the association between alpha oscillatory power and the phenomenological dimension of bliss. Coefficient values and associated metrics are reported, with the formula increasing in model complexity. P-values are uncorrected. SE = Standard Error; BIC = Bayesian Information Criterion.

| **Formula** | **Coefficient** | **SE** | **P-value** | **BIC** |
|---|---|---|---|---|
| Bliss ~ 1 + alpha + (1 \| Subject) | 0.05 | 0.01 | <0.001 | -1389 |
| Bliss ~ 1 + Week + alpha + (1 \| Subject) | 0.03 | 0.01 | 0.034 | -1471 |
| Bliss ~ 1 + alpha + (1 + alpha \| Subject) | 0.04 | 0.11 | 0.71 | -1902 |
| Bliss ~ 1 + Week + alpha + (1 + alpha \| Subject) | 0.02 | 0.11 | 0.85 | -1979 |
| Bliss ~ 1 + Week + alpha + (1 \| Subject:Week) + (1 + alpha \| Subject:Week) | 0.01 | 0.06 | 0.84 | -3689 |
| Bliss ~ 1 + alpha + (1 \| Subject:Week:Session) | -0.02 | 0.01 | 0.047 | -10926 |
| Bliss ~ 1 + Week + alpha + (1 + alpha \| Subject:Week:Session) | 0.04 | 0.03 | 0.23 | -11476 |
| Bliss ~ 1 + alpha + (1 + alpha \| Subject:Week:Session) | 0.04 | 0.03 | 0.21 | -11501 |
| Bliss ~ 1 + Week*alpha + (1 + alpha \| Subject:Week:Session) | 0.02 | 0.06 | 0.71 | -11449 |
| Bliss ~ 1 + Week*alpha + (1 + alpha \| Subject:Week:Session) | 0.02 | 0.06 | 0.71 | -11449 |
| Bliss ~ 1 + Week*alpha + (1 \| Subject:Week:Session) | 0.02 | 0.03 | 0.56 | -10880 |
| Bliss ~ 1 + Condition + alpha + (1 + alpha \| Subject:Week:Session) | 0.07 | 0.03 | 0.007 | -14474 |
| Bliss ~ 1 + Condition*alpha + (1 + alpha \| Subject:Week:Session) | 0.05 | 0.03 | 0.091 | -14456 |

Table S4: A table of the Intraclass Correlation Coefficients (ICCs) of each statistical model within the complexity results are reported here. The ICC was computed by calculating the variance of the random effects of each linear mixed model, and dividing that by the variance of the random effects plus the variance of the residuals of the model. *The statistical model used here was a generalized linear mixed model with a logit link function, with oscillatory alpha power, Global LZc, Global LZSum, and aperiodic exponent included as fixed effects within the model.

| Formula | Variable(s) | ICC |
|---|---|---|
| Data-Driven | Frontal - Occipital (FO) LZSum | 0.57 |
| Data-Driven | Frontal - Frontal (FF) LZSum | 0.56 |
| Data-Driven | Occipital - Occipital (OO) LZSum | 0.57 |
| Data-Driven | Global LZSum (GLZSum) | 0.54 |
| Data-Driven | FO LZc | 0.48 |
| Data-Driven | FF LZc | 0.6 |
| Data-Driven | Global LZc (GLZc) | 0.53 |
| Data-Driven* | Alpha, Exponent, GLZc, GLZSum* | 0.44* |
| Hypothesis-Driven | FO LZSum | 0.55 |
| Hypothesis-Driven | FF LZSum | 0.57 |
| Hypothesis-Driven | OO LZSum | 0.57 |
| Hypothesis-Driven | GLZSum | 0.57 |
| Hypothesis-Driven | FO LZc | 0.53 |
| Hypothesis-Driven | FF LZc | 0.58 |
| Hypothesis-Driven | GLZc | 0.55 |
| Hypothesis-Driven | Alpha, Exponent, GLZc, GLZSum | 0.5 |
| Stimulus-Driven | FO LZSum | 0.7 |
| Stimulus-Driven | FF LZSum | 0.68 |
| Stimulus-Driven | OO LZSum | 0.73 |
| Stimulus-Driven | GLZSum | 0.68 |
| Stimulus-Driven | FO LZc | 0.66 |
| Stimulus-Driven | FF LZc | 0.75 |
| Stimulus-Driven | GLZc | 0.74 |

Table S5: A table of the subject-level intercepts and slopes for a representative data-driven linear mixed model with the formula '(global_lzsum ~ Cluster + (Cluster | Subject) + (Cluster| Subject:Week:Session)'. The asterisks indicate the p-values, which suggest that the participant has a higher/low global complexity value compared to the population average. *p<0.05; ***p<0.001.

| **Subject** | **Intercept** | **SE** | **Slope** | **SE** |
|---|---|---|---|---|
| S01 | -0.17 | 0.22 | 0.05 | 0.1 |
| S02 | 0.13 | 0.2 | -0.14 | 0.09 |
| S03 | -0.07 | 0.22 | -0.06 | 0.09 |
| S04 | 1.01*** | 0.22 | 0.08 | 0.1 |
| S07 | -0.48* | 0.21 | -0.09 | 0.09 |
| S08 | 0.89*** | 0.26 | 0.06 | 0.11 |
| S10 | 0.04 | 0.24 | -0.06 | 0.1 |
| S11 | 0.05 | 0.2 | 0.1 | 0.09 |
| S13 | -0.41 | 0.3 | -0.02 | 0.11 |
| S14 | -0.32 | 0.21 | 0.04 | 0.09 |
| S17 | 0.27 | 0.26 | 0.04 | 0.1 |
| S18 | 0.25 | 0.2 | 0 | 0.09 |
| S19 | -1.12*** | 0.21 | -0.01 | 0.1 |
| S21 | -0.07 | 0.24 | 0.01 | 0.1 |

Table S6:

| **Subject** | **Intercept** | **SE** | **Slope** | **SE** |
|---|---|---|---|---|
| S01 | -0.17 | 0.22 | 0.05 | 0.1 |
| S02 | 0.13 | 0.2 | -0.14 | 0.09 |
| S03 | -0.07 | 0.22 | -0.06 | 0.09 |
| S04 | 1.01*** | 0.22 | 0.08 | 0.1 |
| S07 | -0.48* | 0.21 | -0.09 | 0.09 |
| S08 | 0.89*** | 0.26 | 0.06 | 0.11 |
| S10 | 0.04 | 0.24 | -0.06 | 0.1 |
| S11 | 0.05 | 0.2 | 0.1 | 0.09 |
| S13 | -0.41 | 0.3 | -0.02 | 0.11 |
| S14 | -0.32 | 0.21 | 0.04 | 0.09 |
| S17 | 0.27 | 0.26 | 0.04 | 0.1 |
| S18 | 0.25 | 0.2 | 0 | 0.09 |
| S19 | -1.12*** | 0.21 | -0.01 | 0.1 |
| S21 | -0.07 | 0.24 | 0.01 | 0.1 |
